# Targeting AATF reprograms the tumor microenvironment and suppresses hepatocellular carcinoma via MIR100HG–TGF-β signaling

**DOI:** 10.64898/2026.05.17.725764

**Authors:** Diwakar Suresh, Akshatha N. Srinivas, Bharathwaaj Gunaseelan, S. A. Amith Bharadwaj, Manju Moorthy, Gopalkrishna Ramaswamy, Suchitha Satish, Prashant Vishwanath, Prasanna Kumar Santhekadur, Saravana Babu Chidambaram, Divya P. Kumar

## Abstract

Hepatocellular carcinoma (HCC), a leading cause of cancer death, has a dynamic and heterogeneous tumor microenvironment (TME) that drives progression and therapeutic resistance. We previously elucidated that apoptosis antagonizing transcription factor (AATF) drives angiogenesis in HCC. However, its role in TME remains unexplored. We employed an orthotopic xenograft mouse model, implanting human HCC cells into the liver, and achieved liver-specific silencing via tail vein injection of AAV8 carrying mouse-specific siAATF or siControl. Histological, biochemical, and molecular analyses, combined with whole-genome transcriptomics mapped to mouse and human genomes, were used to study TME and tumor compartments separately. Silencing of AATF in the TME significantly reduced tumor growth compared with controls. Furthermore, AATF loss disrupted key processes in TME, including inflammation, immune response, angiogenesis, and extracellular matrix remodeling. Mechanistically, TGF-β signaling was significantly suppressed in the TME, thereby affecting tumor cell cycle and metabolic activity, ultimately leading to tumor regression. The long noncoding RNA (lncRNA) analysis identified MIR100HG as a key downstream regulator of AATF in the TGF-β signaling pathway. These findings expand the oncogenic role of AATF to include regulation of the TME via the AATF–MIR100HG–TGF-β axis, highlighting its potential as a therapeutic target in HCC.

## Introduction

Hepatocellular carcinoma (HCC) is the primary liver cancer that occurs most frequently, making it the sixth most common malignancy and the third leading cause of cancer-related deaths globally [1]. Its incidence is steadily rising, with projections indicating 1.3 million deaths by 2040 [2]. HCC mainly develops against the backdrop of chronic liver disease, with viral hepatitis and alcohol use being significant risk factors [3]. However, as vaccination programs have significantly decreased hepatitis-related cases, metabolic dysfunction-associated steatotic liver disease (MASLD) and type 2 diabetes have become the main contributors to the increasing global burden of HCC [4]. The rising incidence and mortality rates of HCC present a major healthcare challenge, highlighting the urgent need for heightened attention and intervention [5]. Although there have been advances in HCC treatment, late diagnosis and tumor heterogeneity continue to limit the success of current therapies [6]. Irrespective of etiological factors, persistent hepatic inflammation is a significant risk factor for the development of primary liver carcinoma [7]. About 90% of HCC cases are linked to chronic inflammation that leads to fibrosis, cirrhosis, and ultimately HCC [8]. Epithelial-mesenchymal transition (EMT) and cellular plasticity play essential roles in promoting tumor metastasis [9], posing significant challenges for current molecular-targeted therapies and underscoring the need to discover new disease drivers and treatment options.

The tumor microenvironment (TME), also referred to as tumor stroma, comprises various non-cancerous cells, including fibroblasts, immune cells, endothelial cells, and pericytes [10]. It actively contributes to the regulation of tumor progression through several mechanisms [11]. Chronic inflammation within the TME drives tumor growth by releasing pro-inflammatory cytokines such as IL-6 and TNF-α, while creating an immunosuppressive niche that enables tumor cells to evade immune surveillance through immune checkpoint molecules such as PD-L1/PD-1 [12]. Angiogenesis stimulated by malignant cells secretes vascular endothelial growth factor (VEGF) for a steady supply of oxygen and nutrients necessary for tumor proliferation [13]. Hypoxic conditions within the TME enhance metabolic reprogramming, promote angiogenesis, and increase resistance to apoptosis [14]. Cancer-associated fibroblasts (CAFs) contribute to fibrosis and stromal remodeling by secreting extracellular matrix components, creating a stiff, fibrotic environment that facilitates tumor growth and metastasis [15]. Moreover, metabolic crosstalk between tumor and stromal cells is a dynamic, bidirectional process that promotes the Warburg effect to sustain rapid tumor proliferation [16]. The Tumor microenvironment also induces EMT, enhancing tumor cell motility and invasiveness and facilitating tumor metastasis to other organs [17]. The immune components of the TME, such as myeloid-derived suppressor cells (MDSCs) and CAFs, contribute to chemoresistance by altering drug penetration, enhancing survival signaling, and suppressing immune responses [18, 19]. Together, these factors form a dynamic and adaptable ecosystem that promotes tumor growth, helps cancer evade immune responses, facilitates metastasis, and increases resistance to therapy, highlighting the TME as a key target for treatment. Thus, in contrast to traditional gene therapy that targets genetically unstable tumor cells, recent studies have focused on manipulating the genetically stable tumor microenvironment (TME) to assess its impact on tumor progression [20, 21].

Apoptosis Antagonizing Transcription Factor (AATF) is a coactivator of multiple transcription factors that regulate the cell cycle, DNA damage response, and apoptosis [22–24]. Pathologically, elevated AATF levels have been reported in several cancers, including breast cancer, colorectal cancer, leukemia, multiple myeloma, bladder cancer, and HCC [25–30]. AATF plays a crucial role in tumor proliferation and survival by regulating the G1/S and G2/M cell cycle checkpoints and suppressing the apoptotic response [24,25]. Kumar et al. demonstrated that AATF promotes hepatocarcinogenesis under chronic inflammation conditions induced by MASLD [31]. Previously, we established AATF’s role in regulating angiogenesis in HCC by suppressing pigment epithelium-derived factor (PEDF) [32]. Given its overexpression in HCC, AATF significantly contributes to tumor proliferation and survival. However, despite its recognized oncogenic functions, the role of AATF within the tumor microenvironment (TME) remains unexplored. Understanding how AATF interacts with TME components could reveal additional mechanisms through which it promotes HCC progression.

In this study, we aimed to explore the novel role of AATF in the TME of HCC. Using an orthotopic xenograft mouse model of HCC, we specifically targeted AATF in the tumor stroma or microenvironment (TME) to assess its impact on tumor progression through advanced transcriptomic analyses. Targeting AATF in the TME led to a significant decrease in liver tumor burden. Mechanistically, AATF depletion in TME downregulated TGF-β signaling, subsequently inhibiting extracellular matrix (ECM) remodeling and impairing tumor cell proliferation. Furthermore, we identified the long non-coding RNA MIR100HG as a key regulator in AATF-mediated TME remodeling in HCC. These findings provide new insights into the mechanistic role of AATF in the tumor microenvironment, highlighting its potential as both a malignant biomarker and a promising therapeutic target.

## Results

### Targeting AATF in the tumor microenvironment (TME) inhibits tumor progression in an orthotopic HCC xenograft model

To investigate the potential role of AATF in HCC progression, we established a cell line-derived orthotopic xenograft (CDX) mouse model using human HCC cells in athymic BALB/c (nu/nu) mice. We injected one million cells into the left lateral lobe of the liver through a midline abdominal incision and monitored tumor growth. To achieve liver-specific knockdown of AATF, AAV8 vectors carrying siControl or siAATF under the TBG promoter were administered via tail vein injection. Both groups were observed for four weeks before euthanasia (**Figure 1A**). AATF knockdown in TME significantly reduced tumor growth and invasion compared to the siControl group (**Figure 1B and 1C**). Quantification of tumor weight and volume showed a notable decrease in the siAATF group compared to controls (**Figure 1D and 1E**). Tumor histology revealed fewer poorly differentiated, multinucleated cells with lower mitotic counts in siAATF mice compared to siControl mice, indicating a reduced tumor burden with AATF silencing (**Figure 1F and 1G**).

**Figure 1.**
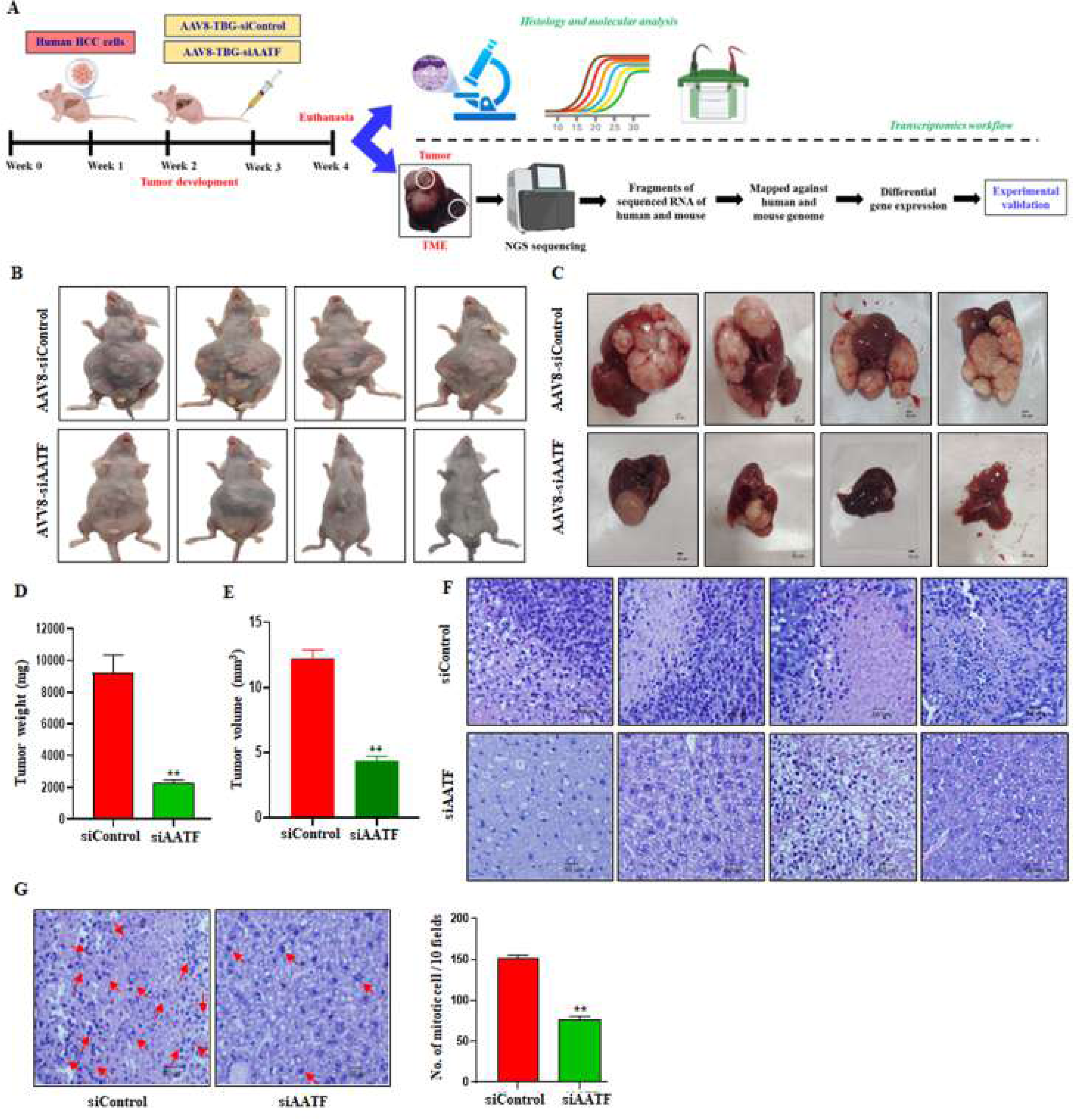
Silencing of AATF reduced tumor burden in the orthotopic xenograft model of HCC. (A) Schematic overview of the study design. Representative images of mice (B) and liver (C) demonstrating tumor formation. Tumor weight (D) and tumor volume (E) were measured. Representative microscopic images of H&E (F) (100x-scale bar 50 μm), and mitotic cell count (G). Data are presented as mean ± SEM for 6-8 mice per group, **p < 0.001 compared to siControl, unpaired t-test. AAV, adeno-associated virus; TBG, tyrosine binding globulin; AATF, apoptosis antagonizing transcription factor; TME, tumor microenvironment.

The liver-specific silencing of AATF was designed to target the tumor microenvironment (mouse cells) surrounding the tumor (human-origin cells). The tumor and TME sections were separated for histological analysis, and immunostaining for AATF confirmed the decreased expression in the TME compared to the tumor sections (**Figure 2A and 2B**). Immunoblotting showed silencing of AATF in the TME of siAATF mice, with no significant changes observed in the tumor tissue or siControl mice (**Figure 2C**). Consistent with the reduced tumor burden, tumor sections were analyzed for Ki-67, a marker of proliferation, and cytokeratin 19 (CK19), a marker associated with aggressive HCC and tumor stemness. AATF silencing resulted in a lower proliferative index and decreased tumor stemness compared to siControl mice (**Figure 2D and 2E**). Additionally, mRNA levels of the HCC marker alpha-fetoprotein (AFP) and the tumor vasculature marker CD31 were notably reduced in the AATF-silenced group, indicating decreased tumor progression and angiogenesis (**Figure 2F and 2G**). Overall, the data demonstrate that silencing AATF within the TME significantly suppresses tumor growth, progression, and invasion.

**Figure 2.**
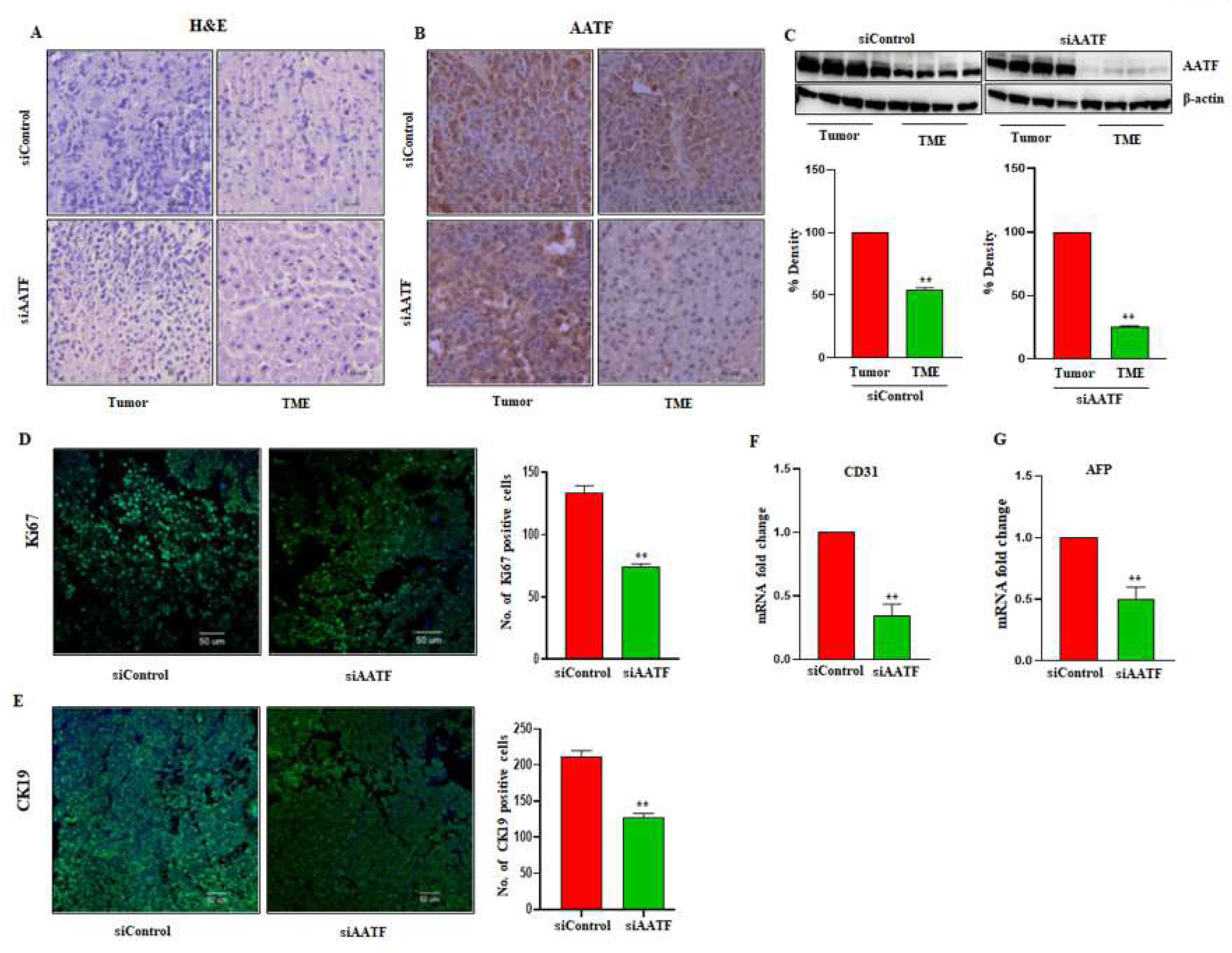
Silencing AATF in TME inhibits HCC tumor progression. Representative microscopic images of (A) H&E staining of tumor and TME sections, (B) immunostaining of AATF (400x), scale bar 50 μm, (C) AATF protein expression in tumor and TME of siControl and siAATF groups. Immunostaining of (D) Ki-67 and (E) CK-19 (200x), Scale bar 50 μm, and mRNA expression of CD31 (F) and AFP (G) were measured in tumor sections of siControl and siAATF mice. Data are presented as mean ± SEM for 6-8 mice per group, **p<0.001 compared to siControl, unpaired t-test. AAV, adeno-associated virus; AATF, apoptosis antagonizing transcription factor; TME, tumor microenvironment; CK19, cytokeratin 19; AFP, alpha-fetoprotein; CD31, cluster differentiation 31.

### Effect of hepatic AATF silencing on transcriptional signatures of the tumor microenvironment and tumor in HCC

To identify the downstream pathways affected by the loss of AATF in the tumor microenvironment (TME), we performed species-specific whole-transcriptomic analysis to distinguish transcriptional changes in the stroma (of mouse origin) from those in the tumor (of human origin). After assessing read quality, the samples were aligned to two reference genomes: human (GRCh38) for tumor cells and mouse (GRCm39) for the TME. Mapping efficiency showed that 87.60% of reads aligned uniquely to the human genome and 60.16% to the mouse genome, highlighting adequate species-specific segregation of transcriptional alterations. Initial dimensionality reduction analysis demonstrated that siAATF and siControl samples formed distinct clusters in both tumor and TME, indicating clear transcriptional separation between the groups and high sample correlation within each group **(Figure S1A-D).** A total of 3943 differentially expressed genes (DEGs) were identified in the TME-specific transcriptome (P-value < 0.05 and | log 2-fold change| ≥ 2). Among these, 1603 genes were upregulated, and 2340 genes were downregulated (**Figure S1E and S1F**). Functional enrichment analysis of DEGs using gene ontology (GO) for biological processes, molecular functions, cellular components, and Reactome pathways revealed that AATF silencing in the TME significantly affected immune-related pathways, angiogenesis, and extracellular matrix remodeling, thereby remodeling the microenvironment to be less supportive of tumor progression (**Figure S1G and Figure S2**). Similarly, 702 genes were differentially expressed in the tumor-specific transcriptome, with 668 upregulated and 34 downregulated (**Figure S1H and S1I**). GO enrichment analysis of the tumor revealed decreased activity in cell cycle, cellular stress response, mismatch repair, DNA damage response, and RNA degradation, resulting in halted cell proliferation (**Figure S1J and Supplementary Figure S3**). Therefore, silencing AATF in the tumor microenvironment leads to significant transcriptomic changes, thereby reducing tumor progression in HCC.

### Targeting AATF reprograms the tumor microenvironment and inhibits tumor progression in HCC

The tumor microenvironment plays a vital role in influencing tumor growth, immune evasion, and resistance to anti-tumor therapies [35]. The TME, which encompasses immune regulation, extracellular matrix remodeling, and angiogenesis, fosters a tumor-permissive environment that is crucial for HCC progression [36]. Since AATF silencing in TME reduced tumor progression, we further examined the molecular signatures of the stroma that may underlie this effect. The GSEA analysis of TME characterization revealed a significant reduction in angiogenesis and an altered immune landscape **(Figure 3A)**. Interestingly, AATF silencing in the TME resulted in the downregulation of key inflammatory and immune pathways, including TNF-α signaling, Il-6/JAK/STAT3 signaling, interferon signaling, TGF-β signaling, and mTORC signaling, while also decreasing hypoxia and metabolic stress in siAATF mice compared to siControl **(Figure 3B and 3C).** Furthermore, we observed downregulation of leukocyte migration, mononuclear differentiation, and suppression of immune response-regulating pathways. Interestingly, chemokines such as CXCL2, CXCL3, and CXCL5, which are primarily involved in recruiting immune cells and myeloid-derived suppressor cells (MDSCs), and thereby contributing to tumor progression [37], were also downregulated. Additionally, the downregulation of CCL2, CCL3, and CCL4, which are involved in the differentiation and infiltration of tumor-associated macrophages (TAMs), supports the notion that AATF silencing disrupts chemokine signaling and hampers the recruitment of tumor-promoting immune cells **(Figure 3D)**. This observation was further supported by decreased chemokine production, cytokine-mediated signaling, and macrophage activation and migration, all of which indicate that AATF silencing broadly suppresses inflammatory responses and TAM production **(Figure 3F, Figure S4A and S4B).** We confirmed the mRNA levels of TNF-α, IL-6, and IL-1β and found that all three inflammatory markers were significantly reduced in siAATF mice compared to siControl **(Figure 3E).** Therefore, AATF silencing reconfigures the TME’s inflammatory environment, making it less conducive to tumor growth and progression.

**Figure 3.**
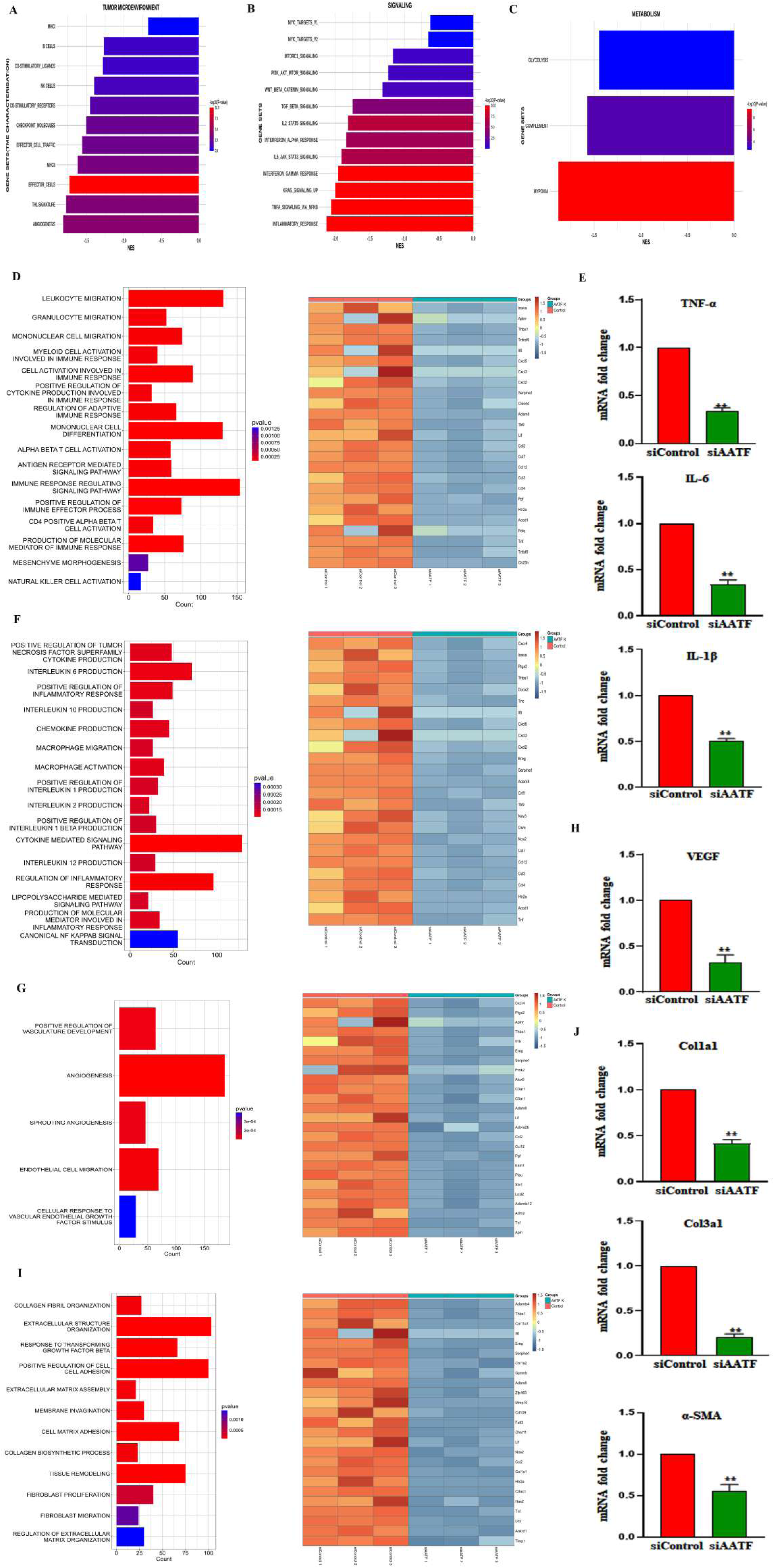
Impact of AATF silencing on key TME components. GSEA analysis of (A) TME characterization, (B) key signaling pathways, and (C) TME metabolism. (D) p-value plots of immune-related pathways and heatmap of the top 25 downregulated genes. (E) mRNA levels of TNF-α, IL-6, and IL-1β. (F) p-value plots of inflammatory pathways and heatmap of the top 25 downregulated genes. (G) p-value plots of angiogenesis pathways and heatmap of the top 25 downregulated genes. (H) mRNA levels of VEGF. (I) p-value plots of ECM pathways and heatmap of the top 25 downregulated genes. (J) mRNA levels of Col1a1, Col3a1, and α-SMA. Data are presented as mean ± SEM for 6-8 mice per group, **p<0.001 compared to siControl, unpaired t-test. TME, tumor microenvironment; TNF-α, tumor necrosis factor-alpha; IL, interleukins; VEGF, vascular endothelial growth factor; Col, collagen; α-SMA, alpha-smooth muscle actin.

Next, we observed that silencing AATF decreased tumor angiogenesis and endothelial cell migration by reducing vascular endothelial growth factor (VEGF) levels (**Figure 3G and 3H**). Since angiogenesis depends on ECM remodeling, particularly the activity of matrix metalloproteinases (MMPs) to clear space for new vessels, we further investigated the ECM-associated components in siAATF mice. Interestingly, extracellular structural organization, tissue remodeling, cell-cell adhesion, and collagen fibril organization were significantly decreased in siAATF mice compared with siControl mice. Notably, the downregulation of LOX, MMPs, TGF-β, α-SMA, Col1a1, and Col3a1 illustrates the signature of ECM destabilization upon AATF silencing **(Figure 3I and 3J).** Therefore, these data strongly suggest that AATF silencing disrupts the structural integrity and remodeling ability of the TME.

TME plays a crucial role in triggering EMT via signaling pathways activated by growth factors, cytokines, and other components of the tumor microenvironment [38]. We examined transcriptional changes to understand how AATF silencing affects EMT-related processes. Our analysis showed that the differentiation and proliferation of epithelial cells were significantly decreased in siAATF mice **(Figure 4A).** While mesenchymal markers and partial EMT markers, such as vimentin, ZEB2, EMP3, and TGF-β, were notably lower in the AATF-silenced group, the epithelial markers PCDH1 and OCLN were significantly higher. These results were further confirmed by q-RTPCR (**Figure 4B and 4C**). Similarly, key EMT-associated pathways such as WNT, Hedgehog, NOTCH, and TGF-β signaling were significantly decreased in siAATF mice compared to controls, along with a low EMT score, as shown by the GSEA of EMT-related genes (**Figure 4D and 4E**). Overall, these findings indicate that AATF silencing inhibits EMT and reduces tumor invasiveness.

**Figure 4.**
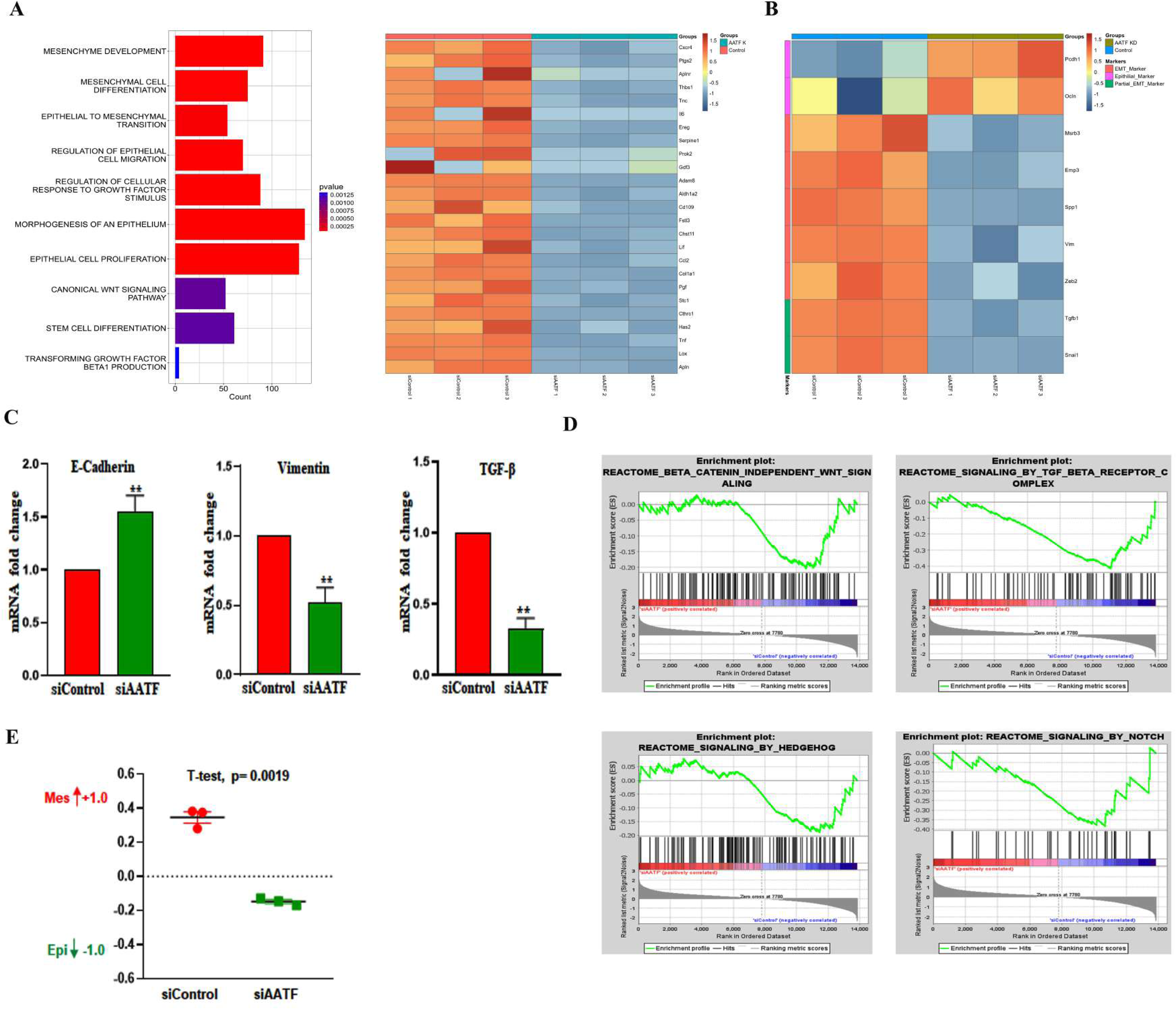
AATF silencing suppresses EMT and decreases tumor invasiveness. (A) p-value plots of downregulated EMT-related pathways and heatmap of the top 25 downregulated genes, (B) Heatmap of epithelial, mesenchymal, and partial EMT markers, (C) mRNA levels of E-cadherin, Vimentin, and TGF-β in siControl and siAATF mice, (D) Enrichment plots for WNT, Hedgehog, NOTCH, and TGF-β signaling pathways, (E) EMT score for siControl and siAATF groups. EMT, epithelial-mesenchymal transition; TGF, transforming growth factor; WNT, wingless-related integration site; NOTCH, neurogenic locus notch homolog.

Next, we examined the compensatory mechanisms that are modulated in the tumor in response to changes in the microenvironment. The main finding from this analysis was that human-origin tumor cells showed reduced cell cycle activity, indicated by the downregulation of CDC16, ECT2, and TCP1, all of which suggest a defective or arrested cell cycle. This implies that AATF silencing has caused cell cycle arrest at multiple points, from cytokinesis to the G1/S checkpoint (**Figure 5A**). Additionally, genes involved in autophagosome maturation, stress kinase activity, and protein K11 ubiquitination were downregulated in tumor cells, indicating metabolic and stress-related dysregulation (**Figure 5B**). Notably, AATF silencing also led to metabolic reprogramming in the tumor cells, as evidenced by decreased sterol synthesis, ATP-dependent activity, and reduced carbohydrate and glucose metabolism (**Figure 5C**). From these findings, AATF silencing in the TME clearly disrupts multiple tumor-supporting mechanisms, including growth arrest, nutrient deprivation, and metabolic shift, leading to tumor regression.

**Figure 5.**
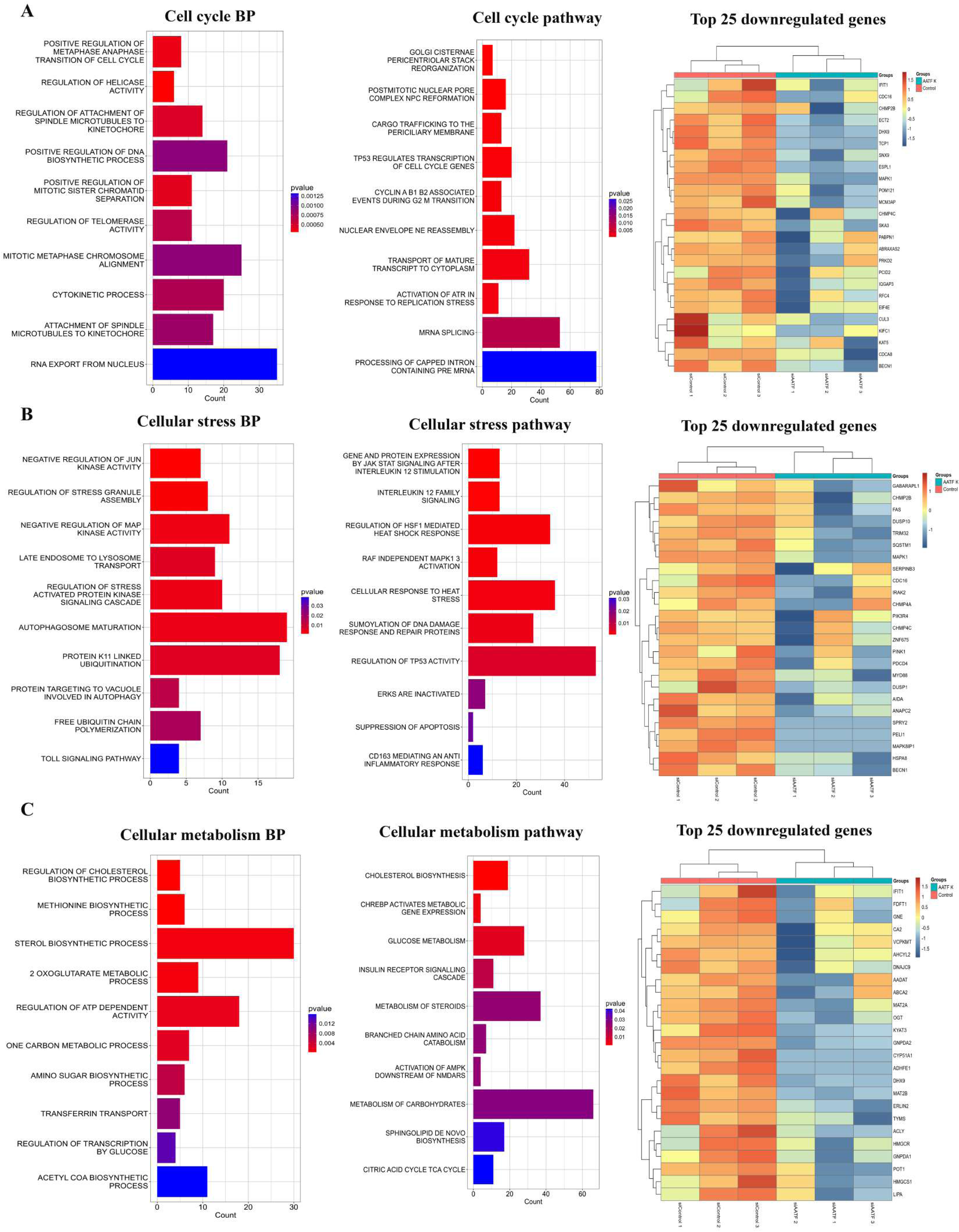
Impact of AATF silencing in TME on tumor growth. (A) p-value plots of downregulated cell cycle-related pathways and heatmap of the top 25 downregulated genes, (B) p-value plots of downregulated cellular stress response-related pathways and heatmap of the top 25 downregulated genes, (C) p-value plots of downregulated cellular metabolism-related pathways and heatmap of the top 25 downregulated genes.

### Targeting AATF in TME suppresses HCC progression via MIR100HG-mediated TGF-β signaling

Since AATF silencing in the TME affected tumor progression and invasion, we investigated tumor-stroma crosstalk using a cell-cell communication explorer (CCCE) [39]. The analysis revealed a significant enrichment of TGF-β signaling from stromal (mouse) cells to tumor (human) cells. Notably, TGF-β–SMAD signaling was markedly reduced in the stromal-to-tumor direction following AATF silencing, indicating disruption of TME-driven pro-tumorigenic signaling (**Figure 6A**). Studies have shown that TGF-β is primarily secreted by cancer-associated fibroblasts (CAFs) in the TME and promotes tumor progression, immune evasion, and metastasis [40–42]. Consistent with this, RNA-seq analysis of the TME compartment demonstrated coordinated downregulation of all three TGF-β ligands (TGF-β1, TGF-β2, and TGF-β3), indicating broad suppression of ligand production. In addition, Tgfbr2 expression was significantly reduced, suggesting impaired receptor-mediated responsiveness within the TME (**Figure S5A-D**). Together, these findings suggest that AATF silencing affects both ligand availability and receptor sensitivity, with a predominant impact on stromal-derived ligand-driven signaling.

**Figure 6.**
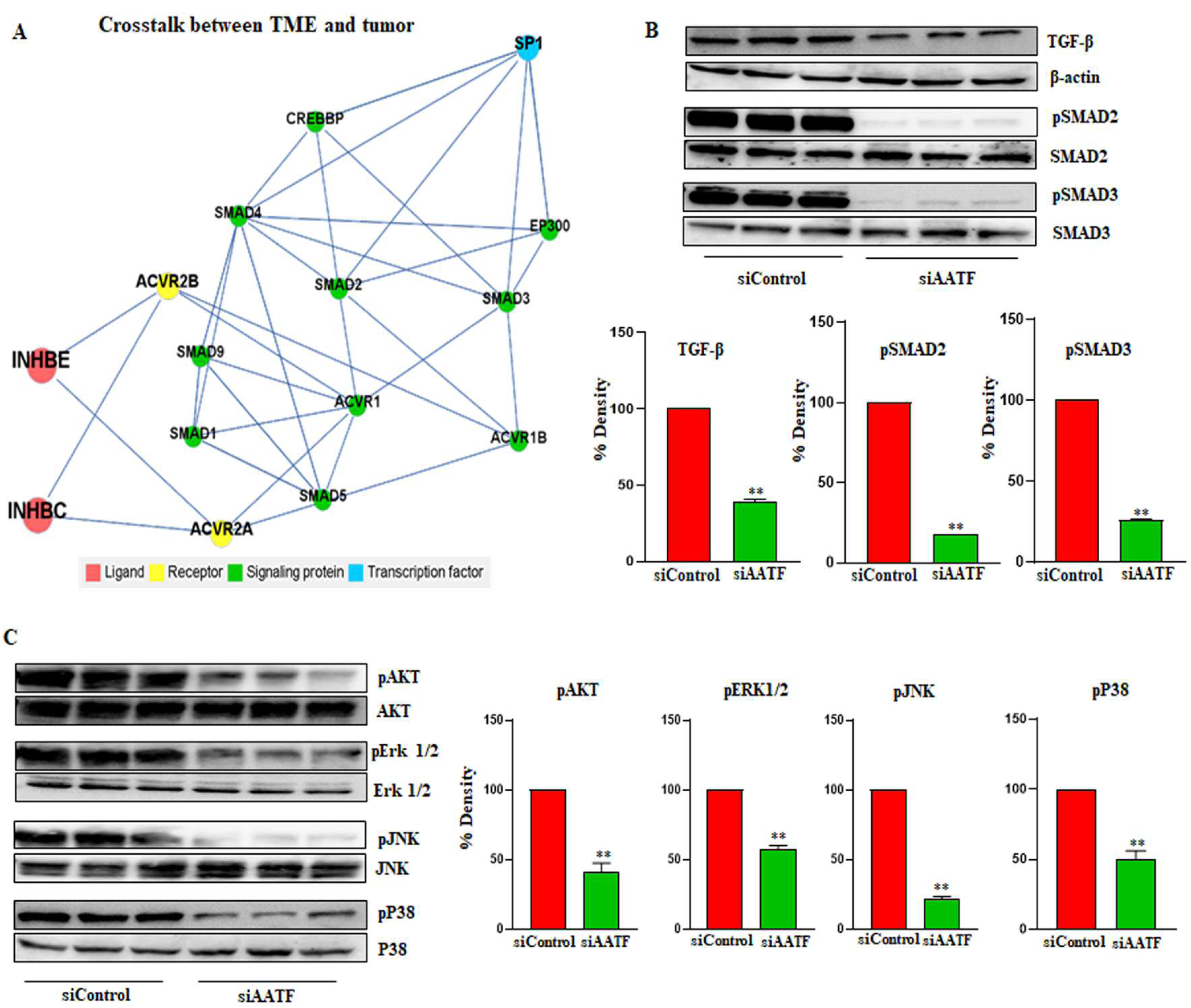
Impact of AATF silencing on tumor-stroma communication. (A) Tumor-TME crosstalk from stromal cells to tumor cells, (B) Western blot analysis of TGF-β, pSMAD2, and pSMAD3, (C) Western blotting of pAKT, pERK1/2, pJNK, and p38MAPK. Data are presented as mean ± SEM for n=3 mice per group, **p<0.001 compared to siControl, unpaired t-test. TME, tumor microenvironment; TGF, transforming growth factor; SMAD, Suppressor of Mothers against Decapentaplegic; AKT, protein kinase B; ERK1/2, extracellular signal-regulated kinase; JNK, Jun N-terminal kinase.

To further validate these observations, we assessed the canonical (SMAD-dependent) and non-canonical (SMAD-independent) pathways of TGF-β signaling. AATF silencing reduced the phosphorylation of SMAD2 and SMAD3, thereby suppressing EMT, immune response, and CAF activation **(Figure 6B)**. Similarly, phosphorylation of AKT and MAP kinases, including ERK, JNK, and p38, was reduced in siAATF mice compared to siControl (**Figure 6C**). Interestingly, MAPK inhibition suppresses cell cycle progression, promotes survival, and facilitates metabolic reprogramming [43]. These data conclusively demonstrate that silencing AATF in the TME reduces TGF-β signaling, thereby affecting tumor-stroma interactions. The decrease in TGF-β signaling within the TME triggers a series of effects that change tumor cell behavior, including decreased cell cycle activity, metabolic reprogramming, and impaired stress response, ultimately leading to reduced tumor growth.

To explore the molecular link between AATF and TGF-β signaling, we examined long noncoding RNAs (lncRNAs) known to enhance or suppress TGF-β expression. Since AATF silencing was specific to the tumor microenvironment (TME), we limited our analysis to mouse-derived long non-coding RNAs (lncRNAs). We quantified known mouse lncRNAs and identified differentially expressed ones using criteria of P-value < 0.05 and |log₂ fold change| ≥ 2. Dimensional reduction analysis revealed distinct clustering between the siAATF and siControl groups, with a strong intra-sample correlation **(Figure S6A-C).** In total, we identified 781 differentially expressed lncRNAs, with 536 upregulated and 245 significantly downregulated in the siAATF group compared to siControl (**Figure S6D and S6E**). Furthermore, to investigate the association between lncRNAs and their direct regulation of target mRNAs, we performed co-expression analysis using the Pearson correlation coefficient. This analysis revealed several co-expressed lncRNA-mRNA pairs, indicating potential regulatory interactions. Functional annotation of the co-expressed mRNAs demonstrated that multiple key pathways were affected by lncRNA-mediated regulation **(Figure S7A)**. Co-expressed lncRNAs with a positive correlation to mRNA downregulation were enriched in GO and Reactome pathways. These pathways align with the TME signature pathways enriched in the mouse transcriptome (**Figure S7B-E**). This indicates that lncRNA-mediated regulation may contribute to the observed immune suppression, ECM remodeling, and reduced angiogenesis, further supporting AATF’s role in shaping a tumor-supportive microenvironment.

Based on the enriched lncRNAs, we built a lncRNA-mRNA pathway interaction network to identify key lncRNAs linked to essential functions of mRNAs (**Figure 7A**). This analysis enabled us to identify critical regulatory lncRNAs that may control ECM remodeling pathways, which are enriched in the network, as downstream effects of AATF silencing in the TME. Among these hub lncRNAs, the top 10 involved in ECM remodeling were further analyzed for functional enrichment to identify lncRNAs that regulate the TGF-β signaling pathway (**Figure 7B**). Among the selected lncRNAs, MIR100HG was significantly downregulated in the siAATF group and showed a strong positive correlation with TGF-β expression. Previous studies have shown that MIR100HG enhances TGF-β signaling by regulating both SMAD-dependent and SMAD-independent pathways [44]. Additionally, MIR100HG encodes miR-100, miR-125b, and miR-99a, which play roles in tumor progression, immune regulation, and metabolic adaptation [45, 46]. Given these findings, we hypothesized that AATF could transcriptionally regulate MIR100HG to activate TGF-β signaling in the TME. Supporting our hypothesis, we observed significant downregulation of MIR100HG in the siAATF mice tissues compared to siControl (**Figure 7C**). To further explore the regulatory mechanism, we performed transcription factor binding site analysis using JASPAR, which revealed strong STAT3 binding sites in the MIR100HG promoter region **(Figure 7D)**. Of note, AATF and STAT3 are transcriptional interacting partners [47]. These findings suggest a potential AATF-STAT3-MIR100HG axis in regulating TGF-β signaling within the TME. Interestingly, phosphorylation of STAT3 was decreased in siAATF mice compared to siControl (**Figure 7E**). Therefore, silencing AATF in the TME decreases MIR100HG levels, which, in turn, affects TGF-β signaling and inhibits tumor growth in HCC **(Figure 7F).**

**Figure 7.**
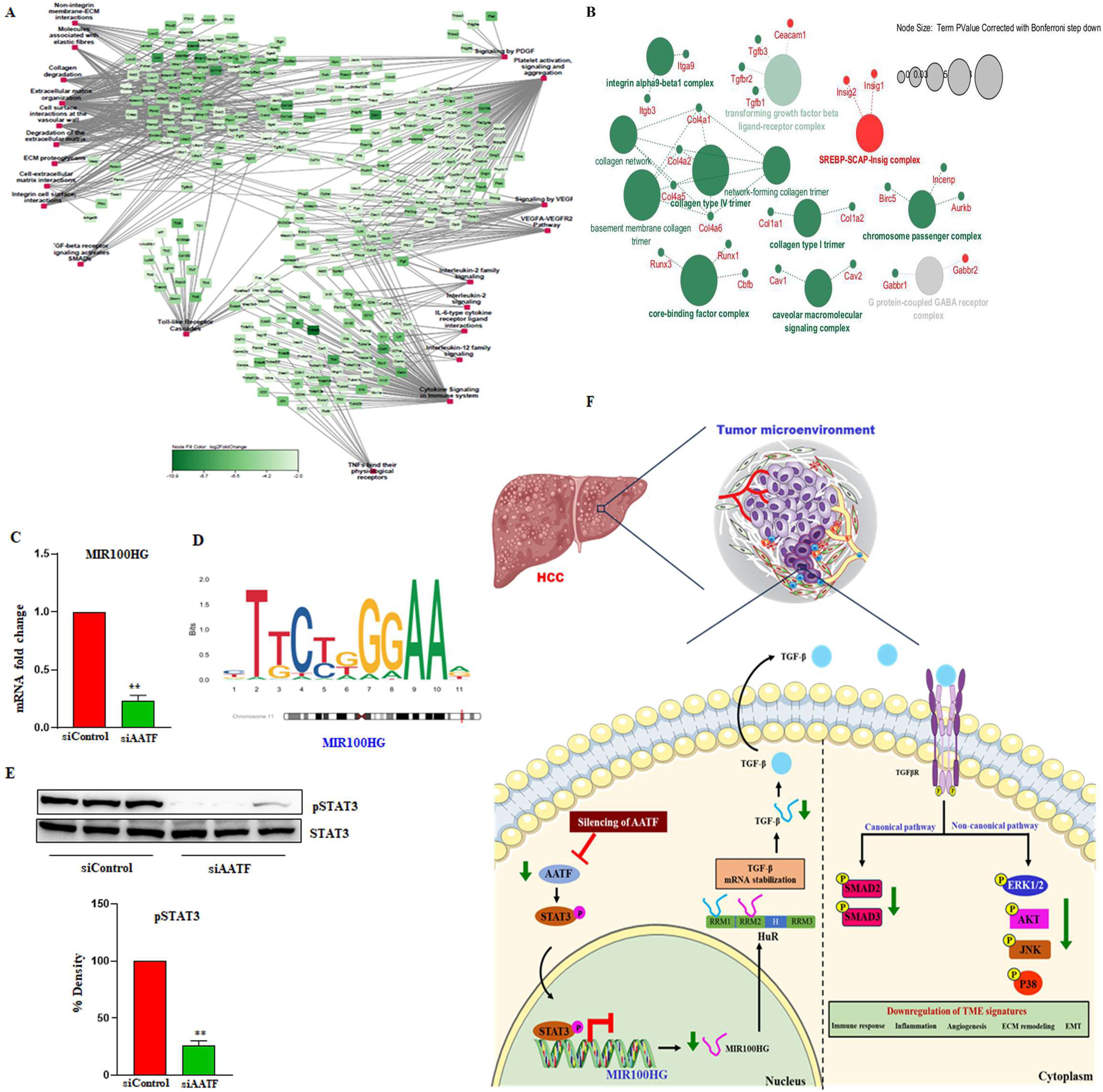
Silencing AATF in TME inhibits TGF-β signaling through MIR100HG. (A) Functional annotation of the top 10 hub lncRNAs (downregulated pathways), (B) Functional annotation of lncRNAs positively co-expressed with TGF-β, (C) qRT-PCR analysis of MIR100HG in siControl and siAATF groups, (D) TFBS analysis using JASPAR for MIR100HG and STAT3, (E) Western blot analysis of pSTAT3 in siControl and siAATF mice, (F) Molecular mechanism illustrating the effect of AATF silencing in the tumor microenvironment and its impact on tumor progression in HCC. lncRNA, long noncoding RNA; TGF, transforming growth factor; qRT-PCR, quantitative real-time polymerase chain reaction; TFBS, transcription factor binding site analysis; STAT3, signal transducer and activator of transcription 3.

## Discussion

HCC is one of the most challenging tumors to treat, mainly because it often has no noticeable symptoms, leading to delayed diagnosis, and due to its varied response to treatment [48]. The complex tumor microenvironment and resistance mechanisms often limit advances in HCC therapy [49]. Emerging therapies, such as dual immunotherapy, antibody-drug conjugates targeting specific markers, and CAR-T cell therapies, have significantly expanded treatment options [50]. However, because these options are less effective, comprehensive efforts to better understand HCC mechanisms could help identify potential disease drivers for improved treatment. The tumor microenvironment, which provides the natural setting, supports tumor cell growth and invasion [51]. This diverse and active population comprises both cellular components (immune cells and other cells that support tumor growth) and non-cellular components (extracellular matrix, cytokines, chemokines, and growth factors), all of which exhibit unique bidirectional communication (52). Tumor cells induce reprogramming of the TME, which, in turn, facilitates tumor heterogeneity and drug resistance [53]. Given the vital role of the TME in cancer progression, targeting it as a treatment has become increasingly common [54]. To understand how tumor-stroma interactions improve clinical outcomes, we explored how AATF influences TME remodeling and promotes tumor growth, and examined unique transcriptional changes in both stromal and tumor regions. Notably, our previous research has demonstrated that AATF promotes angiogenesis in HCC by reducing PEDF levels [32]; however, its role in the tumor microenvironment (TME) remains unclear. Therefore, this study aims to determine whether AATF can function as a regulator of the TME in HCC.

A human HCC cell-derived orthotopic xenograft mouse model was employed to induce tumors in the mouse liver, closely mimicking human disease; the HCC cells (QGY-7703) were authenticated by short tandem repeat (STR) profiling before implantation. AATF was silenced explicitly in the liver using AAV8 vectors encoding a mouse-specific siAATF sequence driven by a convergent promoter. This silencing occurs in the mouse liver, representing the tumor microenvironment (TME) surrounding the implanted human HCC cells. This model closely replicates the clinical scenario of HCC, which develops in a biologically active, often cirrhotic liver microenvironment. Given the well-established roles of liver-derived cytokines, growth factors, and stromal elements in supporting HCC progression [55], our findings prompted us to investigate whether AATF in the hepatic tumor microenvironment contributes to the activation of tumor-promoting signaling pathways. Although siRNA was chosen for gene silencing, it shares shRNA’s stability. The convergent promoter system enables bidirectional expression, a key feature of shRNA, while the AAV8 vector ensures tissue-specific delivery and sustained expression. Our findings demonstrate that silencing AATF in the liver tumor microenvironment (TME) significantly reduced tumor growth, highlighting its role as a key regulator of tumor–stroma interactions. The findings reinforce previous research highlighting the role of stromal factors in tumor behavior [56] and suggest that targeting AATF may interfere with the supportive environment that promotes HCC growth and progression. Additionally, using species-specific transcriptomics (stroma-mouse genes; tumor-human genes, detailed in the Methods section) enabled us to analyze the roles of the stromal and tumor compartments separately, providing a clearer understanding of how AATF affects various cell populations within the TME. The loss of AATF disrupted key processes in the tumor microenvironment (TME), including angiogenesis, extracellular matrix (ECM) remodeling, and immune regulation, ultimately shifting the environment toward an anti-tumorigenic state. Notably, silencing AATF disrupted critical chemokine signals that recruit pro-tumor myeloid cells, such as TAMs and MDSCs, leading to decreased immune cell infiltration and effector activity. Moreover, silencing AATF directly affected inflammatory pathways that promote tumor growth by reducing IL-6, IL-10, and TNF-α levels. Additionally, AATF silencing showed a notable tumor-suppressive effect, evidenced by reduced angiogenesis and endothelial migration. However, the main findings of the study are decreased organization of the extracellular matrix, collagen fibrils, cell-cell adhesion, and tissue remodeling, indicating that AATF silencing significantly impacts the tumor microenvironment (TME) beyond immune suppression and angiogenesis inhibition. Downregulation of ECM proteins such as LOX, MMPs, Col1A1, and Col3A1, along with TGF-β, indicates disruption of ECM integrity, which in turn influences tumor-stroma interactions. AATF silencing also prevented EMT, stopping tumor cells from becoming invasive. The reduction of ZEB2, SNAIL, and Vimentin indicates that tumor cells are less migratory and more epithelial, suggesting that targeting AATF could stop fibroblast-driven ECM remodeling and EMT-induced metastasis. Therefore, by destabilizing the ECM and reducing angiogenic signals, AATF silencing limits both nutrient supply and tumor cell invasion.

Our study showed that silencing AATF in the TME significantly slowed tumor progression, as evidenced by changes in gene expression associated with the cell cycle, stress response, and tumor metabolism. The decreased expression of IFIT1, CDC16, ECT2, and TCP1 indicates a disrupted or paused cell cycle, suggesting that AATF silencing causes cell cycle arrest at various points, including cytokinesis and the G1/S checkpoint. In a pro-tumorigenic environment, stress kinases such as JNK, p38 MAPK, and ERK1/2 are activated by cellular stress, regulating cell survival, apoptosis, and inflammation [57]. Conversely, the downregulation of stress kinase activity in siAATF mice suggests that AATF silencing reduces the tumor’s ability to respond to stress signals, ultimately leading to cell death. Furthermore, the impaired metabolic activity observed upon AATF silencing suggests that tumor cells are metabolically inactive or dormant, potentially leading to tumor regression. Thus, although AATF silencing was confined to stromal cells, human tumor cells exhibited marked transcriptional changes, including cell cycle arrest, suppression of DNA repair and stress responses, and impaired metabolic pathways, highlighting the profound impact of stromal reprogramming on tumor cell behavior.

Our findings highlight the pivotal role of TGF-β signaling in the tumor microenvironment (TME), consistent with its established function in regulating tumor progression and modulating the immune response [58]. Although multiple signaling pathways were altered following AATF silencing, TGF-β signaling was prioritized due to its central role in TME remodeling and its consistent regulation across transcriptomic, signaling, and functional analyses. The altered TGF-β pathway observed upon AATF silencing aligns with reduced cancer-associated fibroblast activation and extracellular matrix remodeling, underscoring AATF’s contribution to maintaining a fibrotic, tumor-supportive niche. The suppression of both canonical (SMAD-dependent) and non-canonical (MAPK, AKT) TGF-β signaling suggests a broad disruption of tumor-stroma crosstalk, which is critical for tumor growth and invasion [59]. Of note, the reduced ECM rigidity and stromal support likely hinder tumor cell invasion and metastasis. Furthermore, downregulation of canonical TGF-β signaling may cause cell cycle arrest by reducing p21 levels. At the same time, non-canonical pathways influence proliferative signaling, highlighting the multiple mechanisms by which AATF silencing impedes tumor progression. These findings position AATF as a key regulator of the pro-tumorigenic microenvironment by modulating TGF-β signaling.

A key finding of our study was the identification of the long noncoding RNA MIR100HG as a downstream mediator of AATF in regulating TGF-β signaling. MIR100HG, which encodes the microRNAs miR-100, miR-125b, and miR-99a, has been linked to controlling both SMAD-dependent and independent TGF-β pathways [60]. Our results demonstrate that silencing AATF reduces MIR100HG expression, thereby decreasing TGF-β signaling and slowing tumor growth. Transcription factor analysis also identified STAT3 as a key regulator of MIR100HG. Given our prior evidence that AATF activates STAT3 transcriptionally, these findings suggest a novel AATF–MIR100HG axis that promotes TGF-β activation within cancer-associated fibroblasts. These findings carry important therapeutic implications. Although TGF-β inhibitors are being explored for HCC treatment, their broad systemic effects have limited clinical efficacy and led to significant side effects. By uncovering AATF as an upstream regulator of MIR100HG-mediated TGF-β signaling, our study suggests a more targeted strategy to modulate the tumor microenvironment without disrupting TGF-β’s essential physiological functions. Targeting the AATF–MIR100HG axis may offer a more selective and effective strategy. Moreover, MIR100HG expression could serve as a biomarker of TGF-β pathway activation, aiding in patient classification for personalized treatments, including combination therapies with immunotherapy.

Our research provides strong evidence that silencing AATF effectively disrupts the tumor-supportive microenvironment, underscoring its potential as a therapeutic target in HCC. Although no specific small-molecule inhibitors of AATF are currently available, indirect strategies, such as disrupting protein–protein interactions, inducing targeted degradation, or modulating TGF-β and STAT3 signaling, may be effective. Future studies using single-cell transcriptomics and spatial approaches will be valuable for identifying specific cellular populations within the TME that are affected by AATF, providing deeper mechanistic insights into immune cell composition and TME remodeling. Although our orthotopic xenograft model effectively demonstrates stromal–tumor interactions, additional research in immune-competent models is necessary to comprehensively understand how AATF affects the immune environment of HCC. Moreover, while MIR100HG appears to be a significant downstream effector, AATF likely impacts other long noncoding RNAs and signaling pathways that warrant further investigation. Primarily, confirming these findings in patient-derived tissues will strengthen their clinical relevance. Moving forward, examining the combined effects of AATF inhibition with TGF-β pathway inhibitors or immune checkpoint therapies could yield new, more effective, and targeted treatments for HCC.

## Conclusion

In this study, we provide compelling evidence for AATF as a stromal regulator in HCC. Our study demonstrates that AATF silencing suppresses tumor progression by remodeling the tumor microenvironment via MIR100HG-mediated modulation of TGF-β signaling. These findings expand the oncogenic repertoire of AATF to include stromal regulation, establish a novel AATF–MIR100HG–TGF-β axis, and reveal a promising therapeutic target for HCC. By integrating transcriptional regulation, lncRNA function, and stromal remodeling, our work provides new mechanistic insights into tumor–stroma interactions and positions AATF as a valuable therapeutic target in HCC.

## Methods

### Orthotopic xenograft mouse model of HCC

Twenty male athymic BALB/c (nu/nu) mice (5 weeks old) were purchased from Hylasco Biotechnology Pvt. Ltd. (Charles River Laboratories, Hyderabad, Telangana). All mice were housed in the central animal housing facility under a 12-hour light-dark cycle with ad libitum access to water and a standard chow diet, and 3–4 mice per cage. The animal study was conducted over 4 weeks, from February 2024 to March 2024, following a 1-week acclimatization period. All animal experiments were approved by the Institutional Animal Ethics Committee of the Center for Experimental Pharmacology and Toxicology (CPT), JSS Medical College, JSS Academy of Higher Education & Research (Approval No. JSSAHER/CPT/IAEC/161/2023), under the supervision of the Committee for the Purpose of Control and Supervision of Experiments on Animals (CPCSEA), Government of India. The experiments were performed in accordance with institutional and national guidelines for the care and use of laboratory animals. All animals were euthanized using a CO₂ chamber following American Veterinary Medical Association (AVMA)-approved humane euthanasia guidelines.

Following one week of acclimatization, the human HCC cells (QGY-7703) cultured at 70-80% confluence were trypsinized, collected, and counted. 1.0 × 10^6 cells suspended in 100 μl of culture medium were mixed with 100 μl of Matrigel (Corning, DC, USA) and orthotopically transplanted into the liver of the mice. The development of the tumor was monitored after tumor inoculation. It was calculated post-euthanasia using the following formula: volume (a rotational ellipsoid) = L × S^2 × 0.5236, where L is the long axis, and S is the short axis.

#### Sample size determination

The sample size was guided by previous liver-specific gene knockdown studies that showed reproducible effects with similar group sizes. Although no formal *a priori* power analysis was performed, 10 mice per group were considered sufficient to detect biologically relevant differences while adhering to the 3Rs principles (Replacement, Reduction, and Refinement).

#### Inclusion and exclusion criteria

All mice that underwent tumor implantation and AAV administration were included in the final analyses. Inclusion criteria required normal postoperative recovery and absence of infection or distress before AAV injection. No animals met exclusion criteria, and no additional exclusion parameters or outlier removal criteria were defined *a priori*.

#### Randomization

Mice were randomly allocated to control and treatment groups. Randomization was performed using a computer-generated random number sequence to assign 10 mice to the AAV8-siControl group and 10 mice to the AAV8-siAATF group.

To minimize potential confounders, mice were housed under identical conditions, and cage positions were randomized within the animal facility to prevent location effects. The order of treatments and measurements was randomized across animals to avoid systematic bias. No other confounders were identified that could have influenced experimental outcomes.

#### Blinding

Group allocation was performed by an investigator not involved in subsequent animal handling or data collection. During the experiment, including AAV administration and measurements, personnel directly handling the animals were aware of group assignments only as required for treatment administration. Outcome assessment and data analysis were performed by investigators blinded to group allocation to minimize observer and analytical bias.

#### Outcome measures

The primary outcome measure was liver AATF expression, which guided sample size selection. Secondary outcome measures included tumor size and molecular markers of inflammation, angiogenesis, and tumor (mRNA and protein levels of relevant target genes). All measurements were performed using validated assays and quantified by investigators blinded to group allocation.

### Liver-specific silencing of AATF with AAV8

Mouse AATF was specifically knocked down in the liver using AAV gene delivery vectors procured from Applied Biological Materials (ABM) Inc., Canada. The siRNA sequences targeting mouse AATF were incorporated into the AAV8 vector under the control of a liver-specific thyroxine-binding globulin (TBG) promoter. The stability of the siRNA was maintained by a convergent promoter vector system that mimics short hairpin RNA (shRNA). Tissue specificity and gene silencing were confirmed using qRT-PCR and Western blotting. One week after tumor implantation, the mice (n=20) were randomly divided into two groups. AAV8-siControl (n=10) and AAV8-siAATF (n=10) were administered via tail vein injection at a dose of 1×10^11^ vg/mouse in a total volume of 200 μl. Mice were monitored for three weeks before euthanasia, and blood and tumor tissues were harvested for further analysis.

### Cell Culture

Human HCC cell line QGY-7703 (CVCL_6715- a kind donation from Dr. Devanand Sarkar, Virginia Commonwealth University, USA) was cultured and maintained in Dulbecco’s modified Eagle’s medium (DMEM) containing 4.5 g/L glucose and supplemented with 10% fetal bovine serum, L-glutamine, and 100 U/ml penicillin-streptomycin at 37°C in 5% CO2. The cell line was authenticated by short tandem repeats (STR) profiling. The experiments were performed with mycoplasma-free cells.

### Tissue processing and analysis

Tumor tissues and the surrounding microenvironmental tissues were isolated, fixed in 4% (v/v) formaldehyde in phosphate-buffered saline, and embedded in standard paraffin blocks. Subsequently, 5-μm-thick tissue sections were cut from each paraffin block and stained with hematoxylin and eosin (H&E). A pathologist at JSS Hospital performed the histopathological analysis of the HCC tissues.

### Immunohistochemistry

The formalin-fixed, paraffin-embedded (FFPE) mouse tissue sections were deparaffinized and rehydrated using xylene and a series of ethanol concentrations. After antigen retrieval with citrate buffer (pH 6) at 94°C for 15 minutes, the sections were incubated with 3% hydrogen peroxide for 10 minutes. Slides were blocked with normal goat serum (Abcam) for 1 hour at room temperature, then incubated overnight at 4°C in a humidified chamber with the AATF antibody (1:500 dilution, Sigma Aldrich). Signals were detected using the Polyexcel HRP/DAB detection system (one-step, PathSitu Biotechnologies) following the manufacturer’s protocol. Nuclear staining was performed with hematoxylin. All immunohistochemistry images were captured using an Olympus BX41 microscope and quantified using ImageJ.

### Immunofluorescence

The FFPE sections were deparaffinized and rehydrated as described above. The slides were incubated with primary antibodies (Ki-67 at 1:250 dilution, Cell Signaling Technologies; CK19 at 1:100 dilution, Santa Cruz Biotechnology), followed by an Alexa Fluor secondary antibody (Thermo Scientific). The nuclei were stained with DAPI (Thermo Scientific). Images were captured with a Leica Stellaris 5 confocal microscope, and analysis was performed in ImageJ.

### RNA isolation and quantitative real-time PCR

Frozen tissues were used to isolate total RNA with TRIzol. RNA was quantified using a Nanodrop spectrophotometer and reverse transcribed into cDNA with the Verso cDNA synthesis kit (Thermo Scientific) following the manufacturer’s protocol. Real-time PCR was performed using the DyNamo Colorflash SYBR Green kit (Thermo Scientific) with 0.5 mM primers (IDT) and 50 ng of cDNA in a 20 μl reaction volume, according to the manufacturer’s instructions on the Rotor-Gene Q5plex HRM System (Qiagen). The relative fold change in mRNA was calculated as 2^(-ΔΔCt) and normalized to the endogenous control β-actin. The validated primer sequences used in this study are provided in Supplementary Table 1.

### Western blotting

The lysates were prepared by homogenizing human liver tissue in 1X RIPA buffer (Sigma Aldrich) containing protease and phosphatase inhibitors (Thermo Scientific). The tissue homogenate was centrifuged at 13,000 rpm for 15 minutes at 4 °C, and the supernatant was collected and stored at -80 °C. The protein concentration was measured using Bradford’s protein estimation method (Bio-Rad). Equal amounts of protein lysates were separated by SDS-PAGE and transferred onto a nitrocellulose membrane for all Western blots. The membranes were blocked with 5% non-fat skim milk for 1 hour at room temperature, then probed with primary antibodies (anti-AATF at 1:1000, Sigma Aldrich; anti-TGF-β, anti-pSMAD2, anti-SMAD2, anti-pSMAD3, anti-SMAD3, anti-pP38MAPK, anti-P38MAPK, anti-pSTAT3, anti-STAT3, anti-β-actin at 1:1000 – Cell Signaling Technologies; anti-pAKT, anti-AKT, anti-pErk1/2, anti-Erk1/2, anti-pJNK, anti-INK at 1:100 – Santa Cruz Biotechnologies) overnight at 4°C. Afterward, the membranes were washed and incubated with secondary antibodies (Cell Signaling Technologies) for 2 hours at room temperature. The blots were developed with the Western Bright ECL HRP substrate (Thermo Scientific), and images were captured using the UVitec Alliance Q9 chemiluminescence imaging system. Band intensities were quantified, and each band was normalized to its respective endogenous control, β-actin.

### Whole transcriptomic analysis

#### RNA isolation, library prep, and sequencing

The RNA was extracted from tissues using the AllPrep DNA/RNA kit (Qiagen) and measured with the TapeStation 4150 and HS RNA screen tape. The library was prepared with the SMART-Seq Library Prep Kit and sequenced with paired-end reads on the Illumina NovaSeq 6000 platform, using a read length of 150 bp × 2.

#### Total RNA-Seq data preprocessing and read alignment

RNA-Seq raw data (FASTQ files) underwent a comprehensive quality assessment using the FastQC program (https://www.bioinformatics.babraham.ac.uk/projects/fastqc/). After the initial QC, low-quality reads were trimmed with Cutadapt v1.15, applying default thresholds except for the minimum read length, which was set at 20 bp. Additionally, preprocessing of total RNA-Seq data and read alignment were performed with a read score threshold of 30. The percentage of reads assigned to the human genome (GRCh38) and mouse genome (GRCm39) was determined using Kraken. Alignment to the reference genome was performed according to previously established protocols [33]. Briefly, a concatenated human (GRCh38) and mouse (GRCm39) genome was constructed as a single reference for alignment with STAR-2.7.11a, allowing up to 3 mismatches per end, discarding reads mapped to multiple locations, removing reads with a mismatch-to-length ratio greater than 0.10, and filtering out noncanonical splice junctions. Remaining parameters adhered to default settings. With these thresholds, reads mapping to both human and mouse genomes were automatically excluded. Gene quantification was performed using species-specific GTF files (Human-gencode.v44.primary_assembly.annotation.gtf, Mouse-gencode.vM33.basic.annotation.gtf) obtained from the Gencode database, utilizing the FeatureCounts program.

#### Differential gene expression analysis

Differential expression analysis was conducted using the Bioconductor package DESeq2 (latest version of R). Genes with read counts fewer than 10 across all samples were excluded before further analysis. Raw read counts were normalized with DESeq2. The magnitude (log2-transformed fold change) and significance (P-value) of differential expression between groups were calculated; genes with P-values < 0.05 were considered significant. We identified DEGs based on the following criteria: P-value < 0.05 and |log2 fold change| ≥ 2. PLS-DA analysis was performed in R using the plsda function in the “mixOmics” package.

#### Functional gene set enrichment analysis (GSEA)

Functional gene set enrichment analysis was performed using the R package ClusterProfiler for Gene Ontology Biological Process (GO BP), Molecular Function (GO MF), and Kyoto Encyclopedia of Genes and Genomes (KEGG) pathways. P-values were adjusted for multiple testing with the Benjamini-Hochberg false discovery rate (BH-FDR). Gene sets with p-values less than 0.05 were considered significantly enriched. These gene sets were downloaded from the MSigDB database for both humans and mice (https://www.gsea-msigdb.org/gsea/msigdb). Additional gene sets related to the mouse tumor microenvironment were obtained from previously published studies [34]. Normalized gene-expression data for the selected pathways were used to generate heatmaps using the pheatmap R package.

#### EMT Scoring

The EMT scoring for each sample was performed using the EMT_Scoring_RNASeq R package (https://github.com/sushimndl/EMT_Scoring_RNASeq). Briefly, raw counts were converted to TPM values, and the EMT score was computed using the 2-sample KS test. A positive EMT score indicates a more mesenchymal phenotype, while a negative score indicates a more epithelial phenotype. In other words, an EMT score closer to +1.0 suggests a more mesenchymal-like (Mes) phenotype, whereas a score closer to -1.0 indicates a more epithelial-like (Epi) phenotype. A Student’s T-test was used to compare the EMT scores of siControl and siAATF, with a P value < 0.05 considered statistically significant.

#### Stroma-Tumor Crosstalk Prediction Using CCC Explorer

The Cell-Cell Communication Explorer (CCC Explorer v.1.1.0) was used to identify tumor (human)–stroma (mouse) crosstalk and associated enriched pathways (P-value < 0.05). DESeq2-normalized data were used as input for the software.

#### Long Noncoding RNA Expression Profiling

Human (gencode.v44.long_noncoding_RNAs.gtf) and mouse (gencode.vM33.long_noncoding_RNAs.gtf) lncRNAs were quantified using Feature Counts and analyzed for differential expression with DESeq2. Raw read counts were normalized by DESeq2. The magnitude (log2-transformed fold change) and significance (P-value) of differential expression between groups were calculated; lncRNAs with P-values <0.05 were considered significant. Differentially expressed lncRNAs with P-value <0.05 and |log2 fold change| ≥ 2 were deemed significant.

To determine the connection between lncRNAs and their directly regulated target mRNAs, we conducted co-expression analysis using Pearson correlation, with a cutoff set to > |0.9| and P value < 0.05. The differentially expressed (log2FoldChange > |2| and P-value < 0.05) mRNA and lncRNA VST-normalized expression data served as input for this analysis. Additionally, FEELnc (FlExible Extraction of LncRNAs) was used to classify co-expressed targets as cis or trans. mRNAs that were either positively or negatively co-expressed were analyzed for their enriched functions using Gene Ontology and Reactome pathways through the ClusterProfiler package. In addition to the standard mRNA-target-based functional prediction, we also used the Ncpath database (http://ncpath.pianlab.cn) to identify enriched KEGG pathways among the significantly differentially expressed long non-coding RNAs (lncRNAs) in humans.

Furthermore, for pathways enriched for lncRNA targets, a lncRNA-mRNA-pathway network was constructed in Cytoscape v.3.10.0. Because many lncRNAs co-express with the selected mRNA, they were displayed side by side as heatmaps, and these plots were added to the network using the enhanced graphics application.

The top 10 hub lncRNAs, which target the most mRNAs, were used to build a coding-non-coding gene co-expression network in Cytoscape v.3.10.0. To predict the functions of the hub lncRNA targets, the Cytoscape plug-in ClueGo was employed. Since it was found that the number of mRNA targets of the mouse hub lncRNA was large, only the top 10 positively and negatively correlated mRNA targets of the hub lncRNA were used as input for the co-expression network formation.

#### Analysis of Transcription Factor Binding Sites

We analyzed predicted TF binding motifs in the 2-kb promoter region (and an additional 100 bp downstream) of the human lncRNA MIR100HG using the JASPAR2024 database, prioritizing JASPAR scores above 400. The sequence logo was generated using the R package TFBSTools and was used to query the STAT3 motif (ID: MA0144.2). The GGseqlogo package was used to plot the sequence logo for STAT3. Furthermore, the MIR100HG promoter was visualized using the Gviz package in R to plot the MIR100HG promoter region. The GenomeAxisTrack and IdeogramTrack functions were used to visualize the chromosome and its corresponding start and end positions.

### Statistical analysis

The data were expressed as mean ± SEM. Statistical significance between the two groups was determined using Student’s t-test. All statistical analyses were conducted with GraphPad Prism software (version 8), and p-values < 0.05 (*) or < 0.001 (**) were regarded as significant. P < 0.05 was considered to indicate a statistically significant difference. All statistical analyses and visualizations for whole-transcriptomic data were performed in R version 4.3.2, unless otherwise specified.

## Acknowledgments

All animal experiments were approved by the Institutional Animal Ethics Committee of the Center for Experimental Pharmacology and Toxicology (CPT), JSS Medical College, JSS Academy of Higher Education & Research (Approval No. JSSAHER/CPT/IAEC/161/2023), under the supervision of the Committee for the Purpose of Control and Supervision of Experiments on Animals (CPCSEA), Government of India.

This study was fully or partially supported by the Extramural Ad-hoc Grant from the Indian Council of Medical Research (ICMR-Grant No.: 5/3/8/55/2020-ITR) awarded to DPK, and a Research Fellowship from JSS AHER granted to DS.

RNA sequencing was performed at Molsys Pvt. Ltd. (Molsys Scientific). We thank TheraCUES Innovations Pvt Ltd. for the assistance with the bioinformatic analysis of the whole transcriptomic data. We also thank the confocal instrumentation facility (Leica Confocal Microscope Stellaris 5, funded by DST-PURSE, SR/PURSE/2021/81-(c) year 2022) at the University Sophisticated Instrumentation Center (USIC), JSS AHER.

The authors also acknowledge funding support from the Department of Biotechnology—Boost to University Interdisciplinary Life Science Departments for Education and Research program [DBT-BUILDER: BT/INF/22/SP43045/2021] and the Department of Science and Technology–Promotion of University Research and Scientific Excellence [DST-PURSE: SR/PURSE/2021/81(c)].

## Authors’ contributions

DS: Investigation, Data curation and analysis, Writing-original draft, review and editing; ANS: Data curation and analysis, Writing-review and editing; BG and AB: Methodology, Investigation, Writing-review and editing; MM and GR: Data analysis, Writing-review and editing; SS: Investigation, data analysis, Writing-review and editing; PV, PKS, and SBC: Conceptualization, Writing-review and editing; DPK: Conceptualization, Investigation, Validation, Supervision, Data analysis, Funding acquisition, Writing-original draft, review and editing. All authors have read and approved the final version of the manuscript.

## Availability of data and materials

Data presented herein are available from the corresponding author upon request. Whole genome sequencing data have been uploaded to the GEO database (Accession number: GSE295864).

---

**Figure S1.**
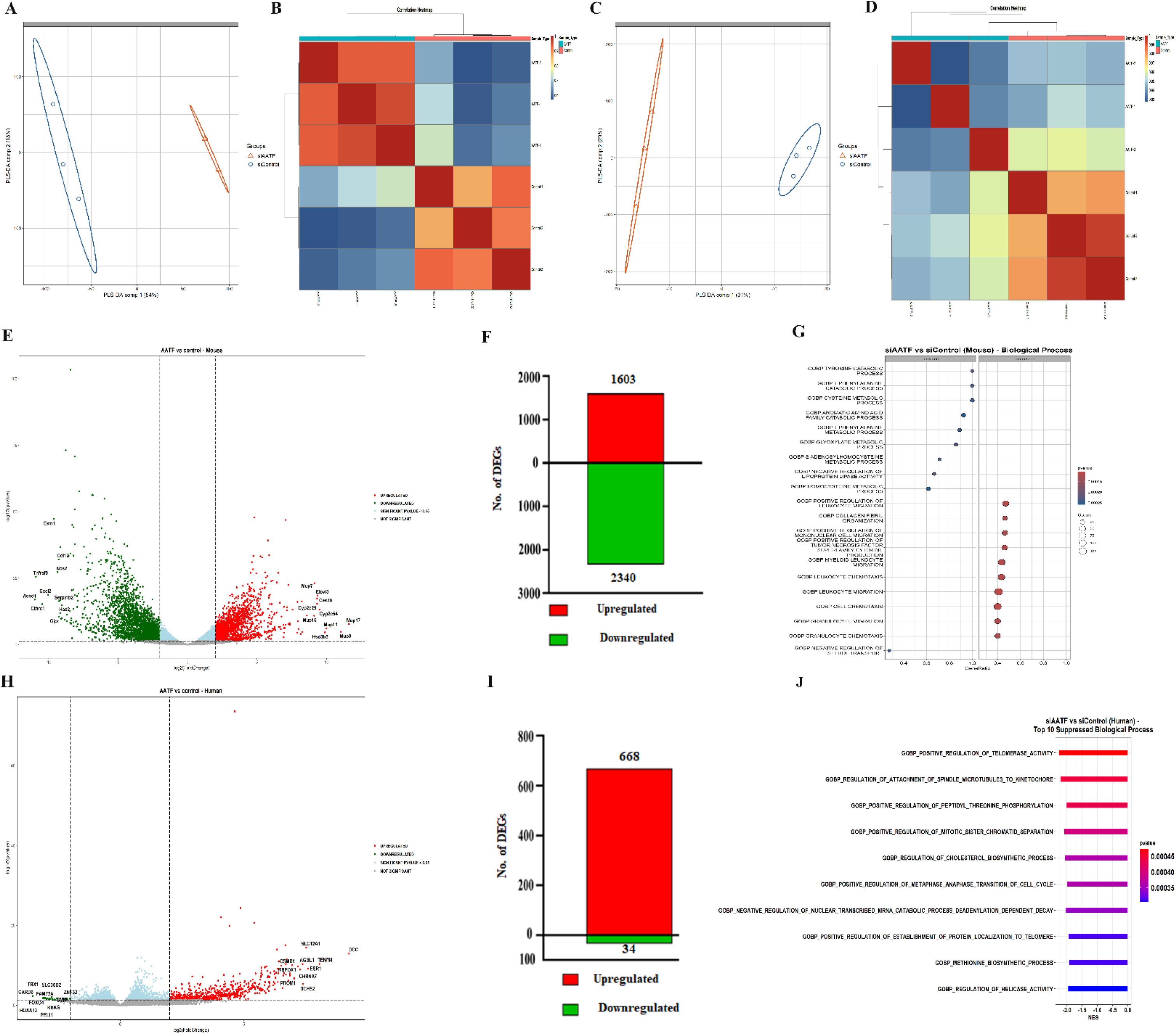
Differential gene expression in tumor and TME following AAV8-TBG-siAATF treatment in an orthotopic xenograft model of HCC. (A) PLS-DA plot, (B) sample correlation plot for TME, (C) PLS-DA plot, and (D) sample correlation plot for tumor showing clear separation between groups, (E) Volcano plot for TME displaying differentially expressed genes. Red dots indicate gene upregulation, green dots indicate downregulation, and grey dots are unchanged (p<0.05, log fold≥ 2). (F) Number of genes up- and downregulated in TME transcriptome, (G) Top 10 up- and downregulated GO-BP in siAATF versus siControl group in TME, (H) Volcano plot for tumor transcriptome illustrating DEGs with red dots for upregulation, green for downregulation, and grey dots unchanged (p<0.05, log fold≥ 2). (I) Number of genes up- and downregulated in tumor transcriptome, (J) Top 10 downregulated GO-BP in siAATF versus siControl group in tumor. AAV, adeno-associated virus; AATF, apoptosis antagonizing transcription factor; TME, tumor microenvironment; PLS-DA, partial least squares discriminant analysis; GO, gene ontology; BP, biological processes.

**Figure S2.**
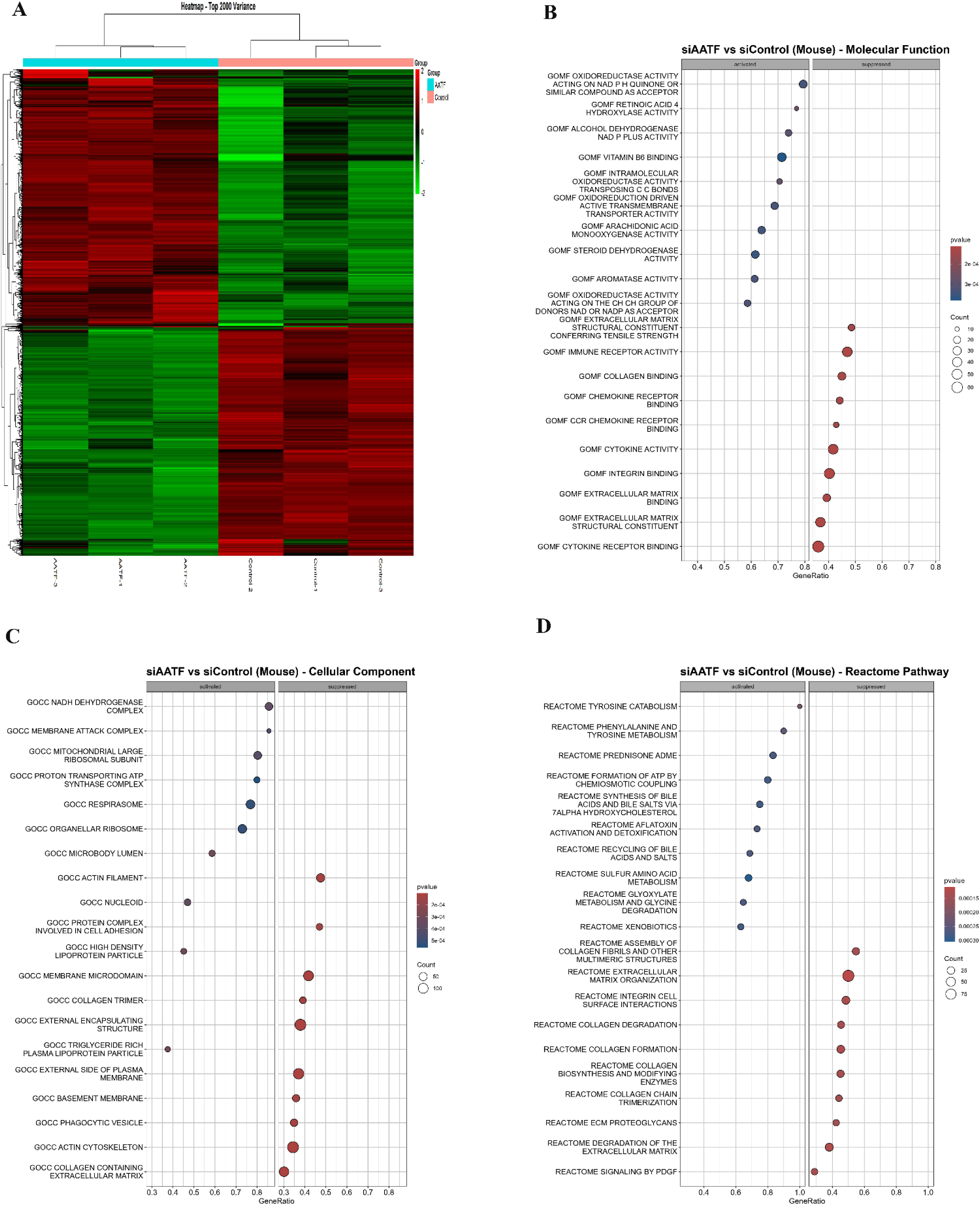
Hierarchical clustering and pathway enrichment analysis of <u>TME</u> in siAATF versus siControl groups. (A) Hierarchical clustering of siAATF versus siControl groups in TME. The top 10 upregulated and downregulated terms in (B) GO-MF, (C) GO-CC, and (D) Reactome pathways in siAATF versus siControl groups in TME. GO, gene ontology; MF, molecular function; CC, cellular component.

**Figure S3.**
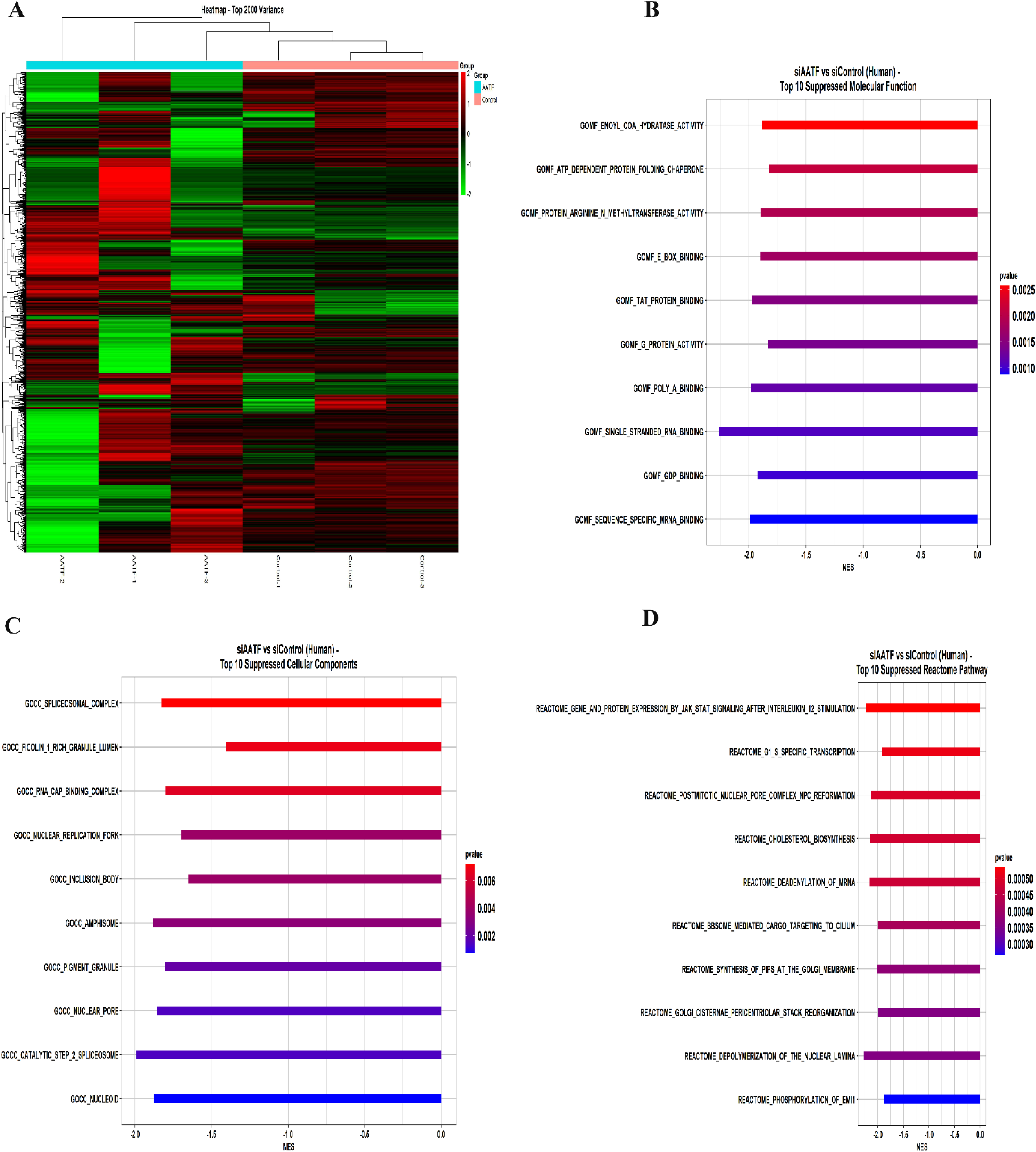
Hierarchical clustering and pathway enrichment analysis of <u>tumors</u> in siAATF versus siControl groups. (A) Hierarchical clustering of siAATF versus siControl groups in the tumor. The top 10 downregulated terms in (B) GO-MF, (C) GO-CC, and (D) Reactome pathways in siAATF versus siControl groups in TME. GO, gene ontology; MF, molecular function; CC, cellular component.

**Figure S4.**
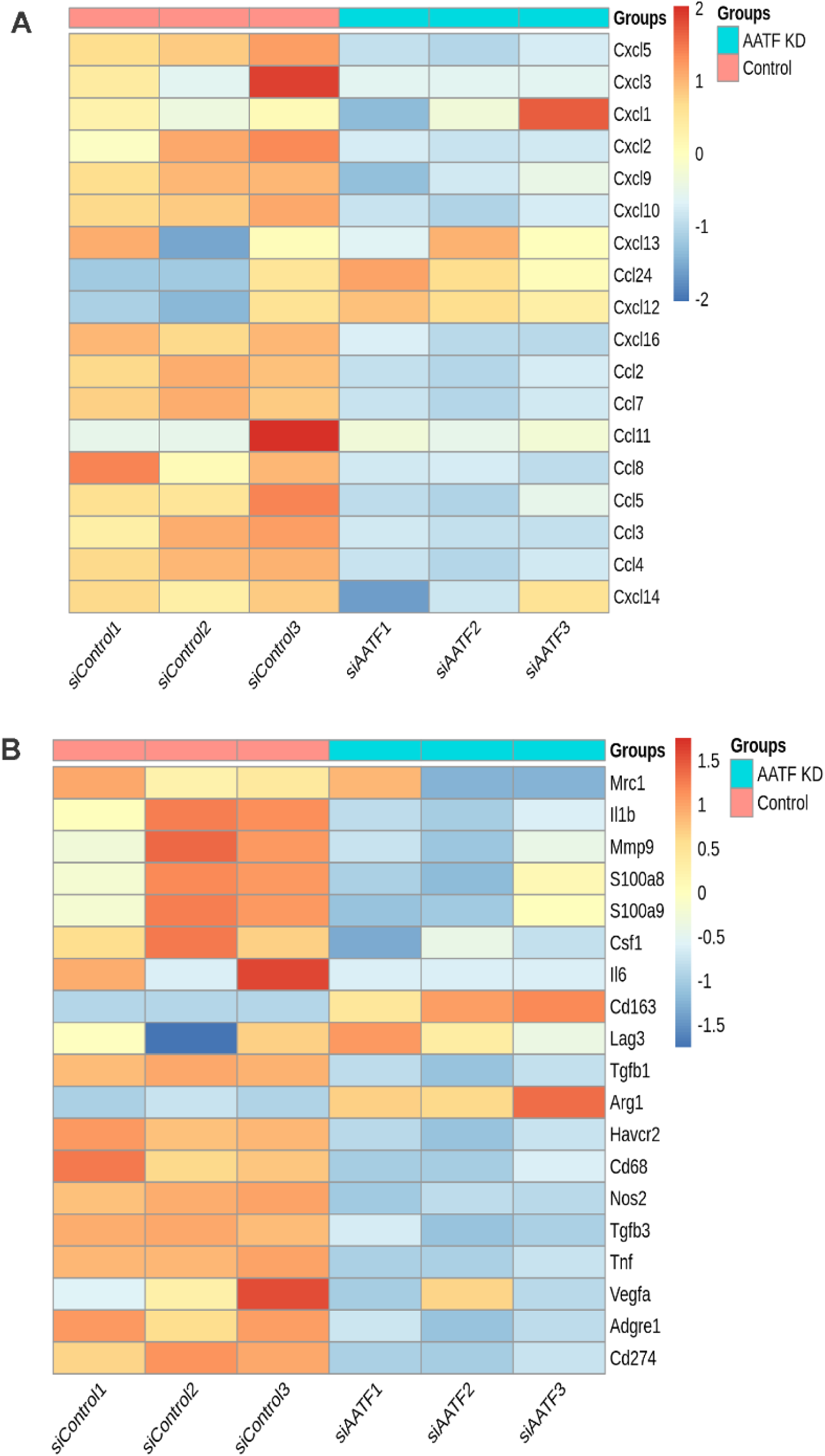
Downregulation of chemokine and immune remodeling genes upon AATF silencing in the TME. Heatmap of chemokines (A) and immune remodeling (B) genes in siAATF versus siControl groups in TME.

**Figure S5.**
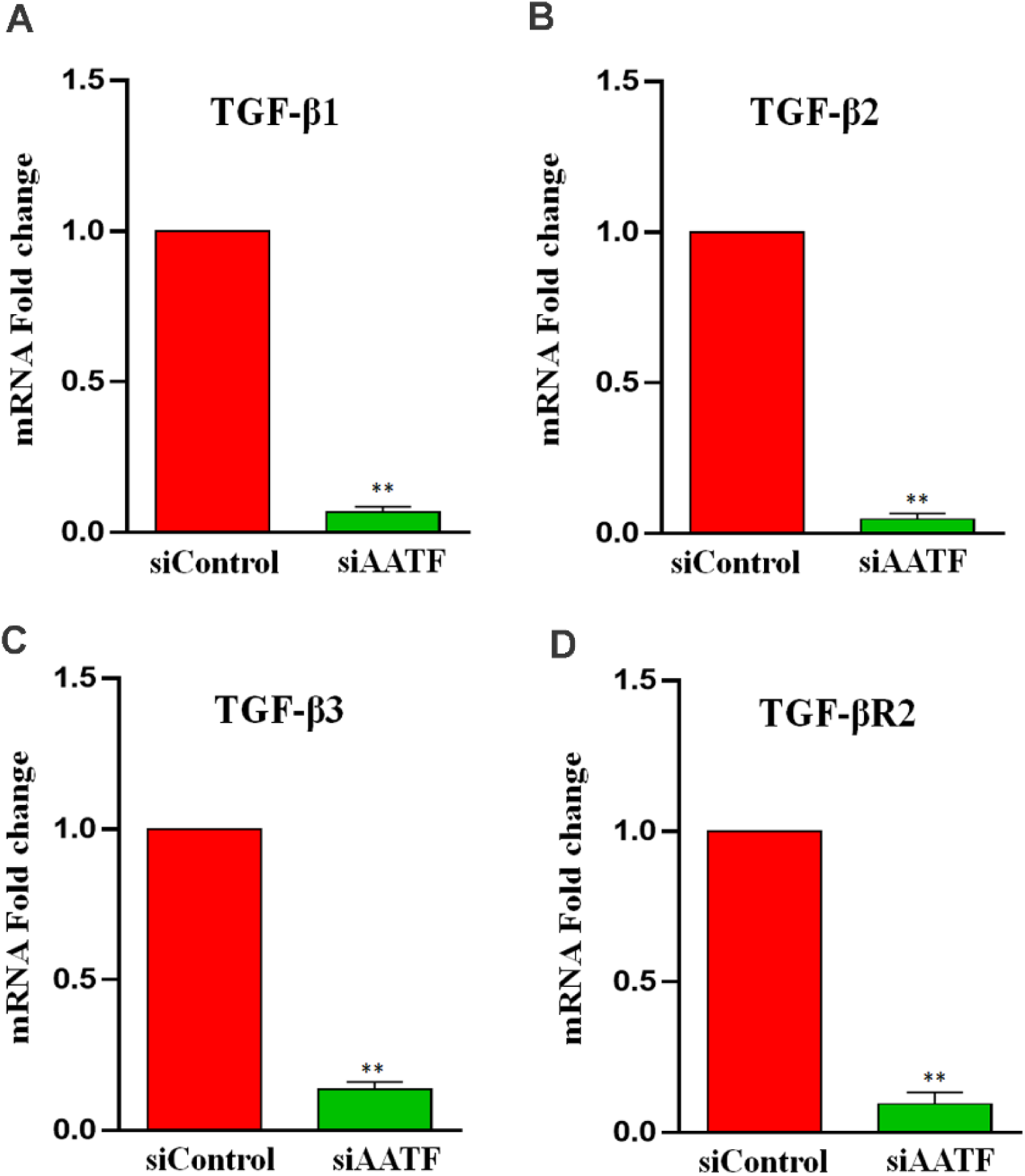
Expression profiling of TGF-β ligands and receptors. mRNA expression of TGF-β1 (A), TGF-β2 (B), TGF-β3 (C), and TGF-βR21 (D) expressed as fold change. Data are presented as mean ± SEM for n=3 per group, **p<0.001 compared to siControl, unpaired t-test. TGF-β, transforming growth factor-beta; TGF-βR, transforming growth factor-beta receptor.

**Figure S6.**
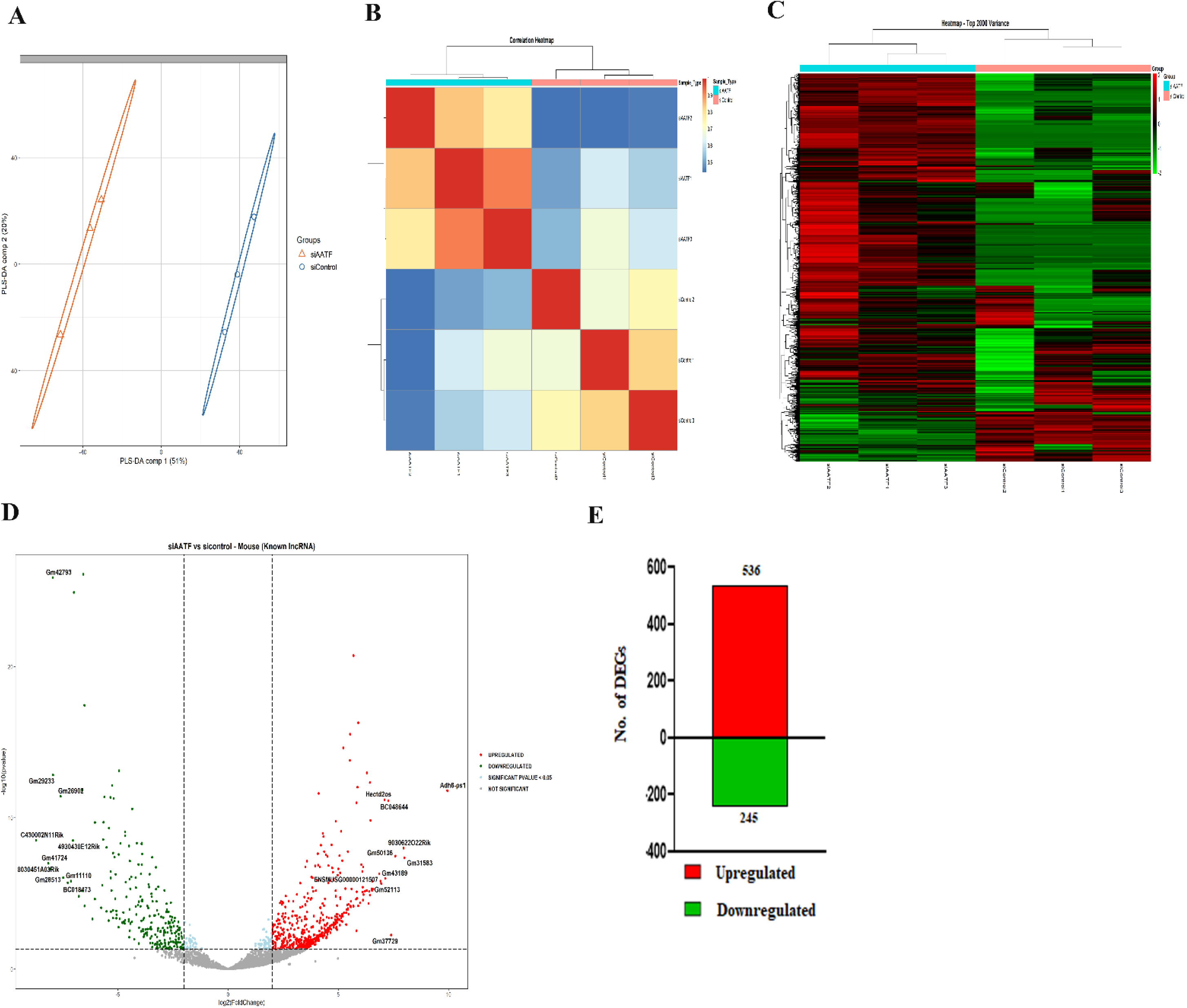
Transcriptomic analysis of mouse lncRNAs in the tumor microenvironment. (A) PLS-DA plot, (B) sample correlation plot, (C) hierarchical clustering of mouse lncRNAs, (D) Volcano plot showing differentially expressed lncRNAs. Red dots indicate gene upregulation, green dots indicate gene downregulation, and grey dots are unchanged (p<0.05, log fold≥ 2), and (E) number of lncRNAs upregulated and downregulated in TME.

**Figure S7.**
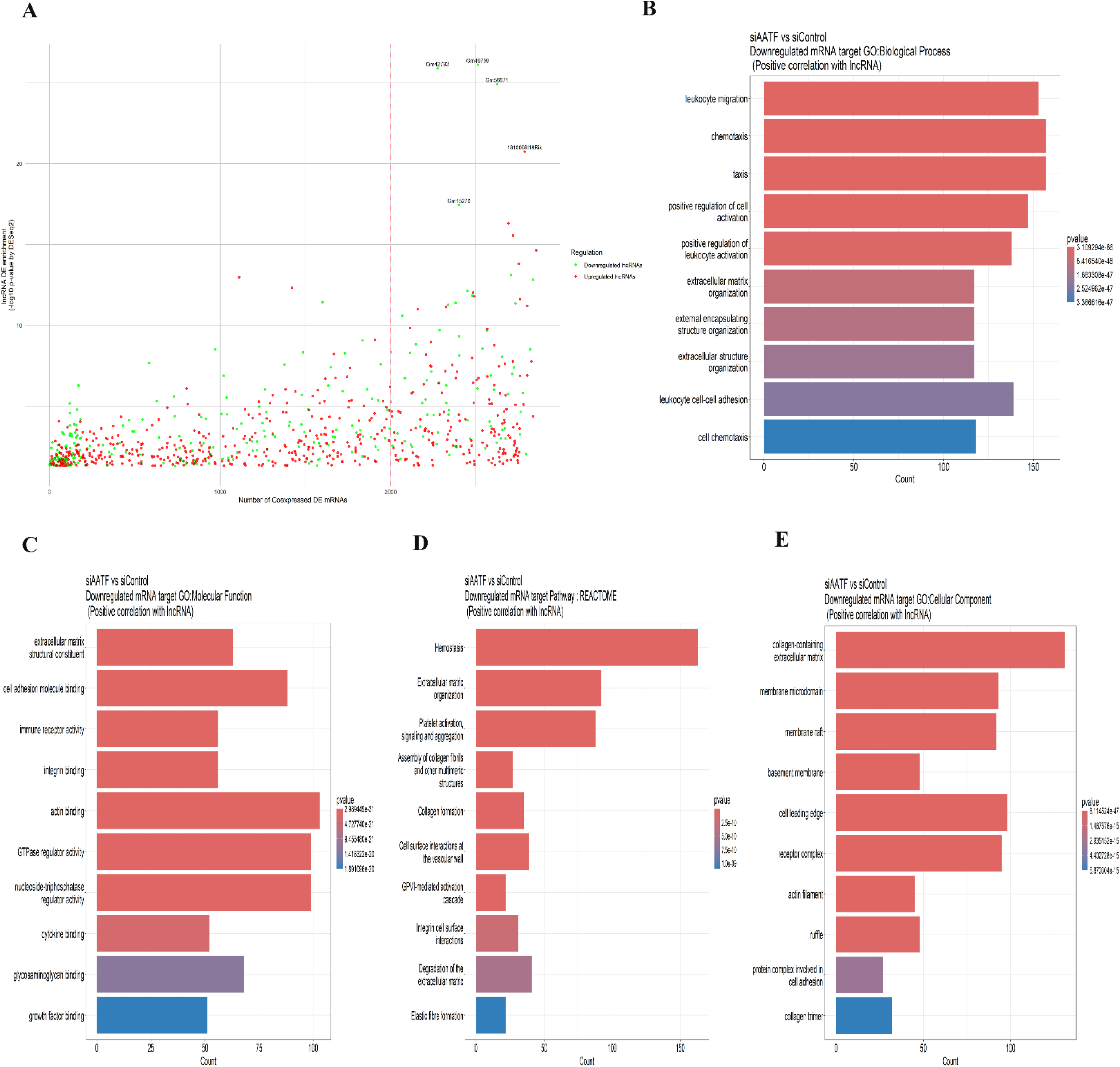
Correlation and functional enrichment analysis of downregulated lncRNA and mRNA. (A) Pearson’s correlation analysis of co-expressed lncRNA and mRNA, (B) GO-BP, (C) GO-MF, (D) GO-CC, and (E) Reactome pathway analysis for positively correlated downregulated lncRNA-mRNA.

**Table S1.**
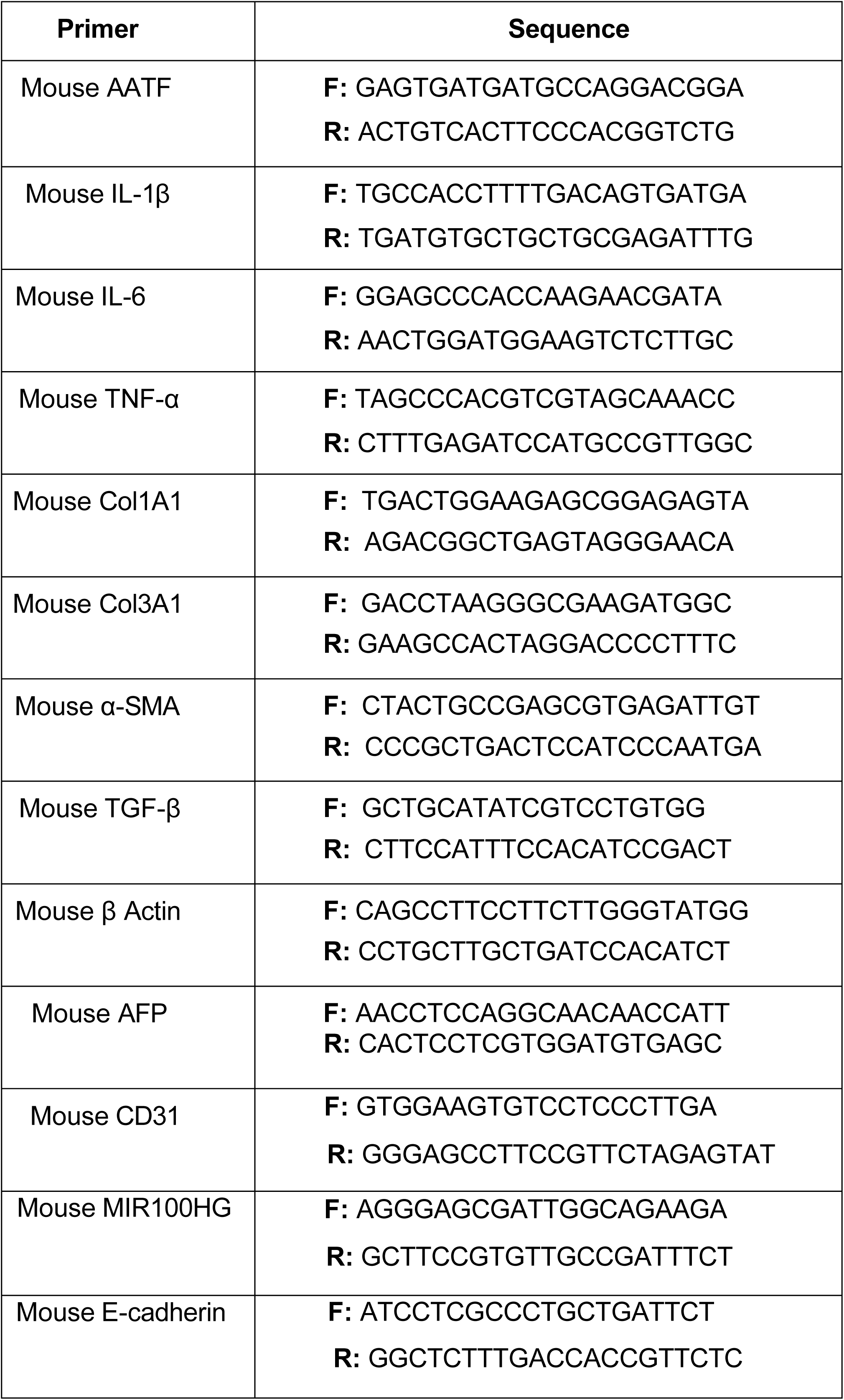
List of primer sequences used in qRT-PCR in the study.

